# Evolution of the potassium channel gene *Kcnj13* underlies colour pattern diversification in *Danio* fish

**DOI:** 10.1101/2020.06.25.170498

**Authors:** Marco Podobnik, Hans Georg Frohnhöfer, Christopher M. Dooley, Anastasia Eskova, Christiane Nüsslein-Volhard, Uwe Irion

**Author notes:** Corresponding author Correspondence to Uwe Irion.

## Abstract

The genetic basis of morphological variation provides a major topic in evolutionary biology^1-6^. Colour patterns in fish are among the most diverse of all vertebrates. Species of the genus *Danio* display strikingly different colour patterns ranging from horizontal stripes, to vertical bars or spots^7-10^. Stripe formation in zebrafish, *Danio rerio*, oriented by the horizontal myoseptum, is a self-organizing process based on cell-contact-mediated interactions between three types of chromatophores with a leading role of iridophores^11-14^. We investigated genes known to regulate chromatophore interactions in zebrafish as candidates that might have evolved to produce a pattern of vertical bars in its sibling species, *Danio aesculapii*^8,10^. Using gene editing^15-17^ we generated several mutants in *D. aesculapii* that demonstrate a lower complexity in the interactions between chromatophores in this species, as well as a minor role of iridophores in patterning. Complementation tests in interspecific hybrids^18,19^ identified *obelix/Kcnj13*, which encodes an inwardly rectifying potassium channel (Kir7.1)^20^, as a gene evolved between *D. rerio* and *D. aesculapii* as well as in two of seven more *Danio* species tested. Our results demonstrate that the CRISPR/Cas9-system allows straightforward genetic tests also in non-model vertebrates to identify genes that underlie morphological evolution.

## Main

Colour patterns are common features of animals and have important functions in camouflage, as signals for kin recognition, or mate choice. As targets for natural and sexual selection, they are of high evolutionary significance^21-24^. The zebrafish, *Danio rerio*, has emerged as a model system to study colour pattern development in a vertebrate^14,25-29^. A fair number of genes have been identified that are required for the formation of the pattern^28,29^, which is composed of a series of horizontal light and dark stripes on the flank of the fish as well as in the anal and tail fins (Fig. 1a). The adult pattern is created by three different types of pigment cells (chromatophores) in the skin, black melanophores, blue or silvery iridophores and yellow xanthophores^13,30-32^. The chromatophores producing this pattern mainly originate from multipotent neural crest–derived stem cells located at the dorsal root ganglia of the peripheral nervous system^33-37^. Signalling pathways, e.g. Csf1 or Edn3, control proliferation and spreading of chromatophores^38-40^. During metamorphosis, the period when adult form and colour pattern are established, assembly into the striped pattern is controlled by interactions between the three cell types. Several genes are autonomously required in the chromatophores for these heterotypic interactions^11,12,20,41-43^. These genes typically encode integral membrane proteins such as adhesion molecules^42^, channels^20^, or components of cellular junctions, some of which mediate direct cell contacts^43,44^. In *Meox1* (*choker*) mutants, lacking the horizontal myoseptum as anatomical landmark, the horizontal orientation is lost, but stripes form of normal width and composition (Fig. 1b), indicating that stripe formation is a process of self-organization of the pigment cells^11^.

**Fig. 1:**
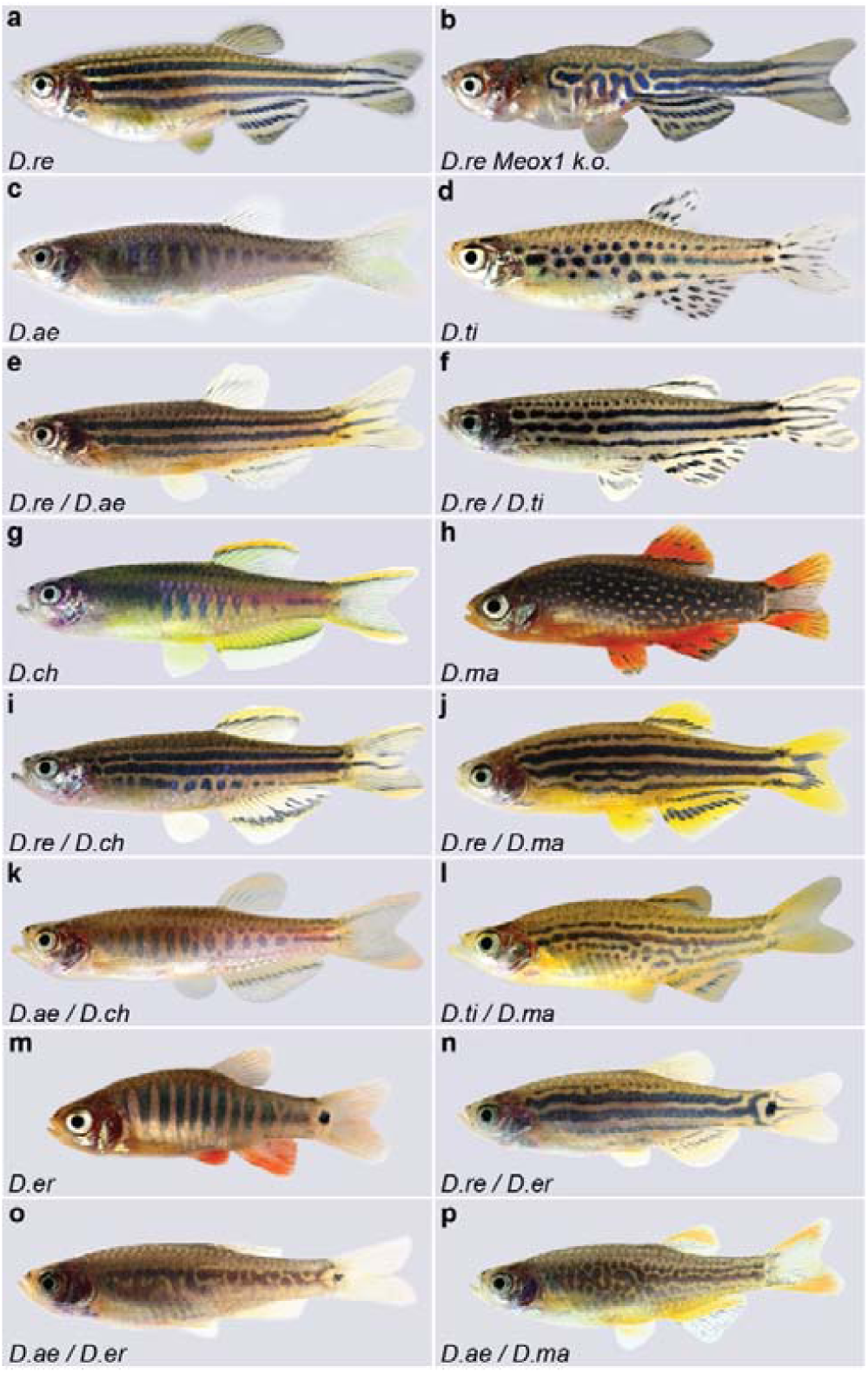
Colour patterns in *Danio* fish and interspecific hybrids. **a**, Colour pattern of zebrafish, *D. rerio* (*D*.*re*). **b**, *D. rerio Meox1* (*choker*) mutants, which lack a horizontal myoseptum. **c**, *D. aesculapii* (*D*.*ae*). **d**, *D. tinwini* (*D*.*ti*). **e**, Interspecific hybrid between *D. rerio* and *D. aesculapii* and, **f**, between *D. rerio* and *tinwini*. **g**, *D. choprae* (*D*.*ch*). **h**, *D. margaritatus* (*D*.*ma*). **i** Interspecific hybrid between *D. rerio* and *D. choprae*, **j**, and between *D. rerio* and *D. margaritatus*. **k**, Interspecific hybrid between *D. aesculapii* and *D. choprae*. **l**, Interspecific hybrid between *D. tinwini* and *D. margaritatus*. **m**, *D. erythromicron (D. er)*. **n**, Interspecific hybrid between *D. rerio* and *D. erythromicron*. **o**, Interspecific hybrid between *D. aesculapii* and *D. erythromicron*. **p**, Interspecific hybrid between *D. aesculapii* and *D. margaritatus*.

*Danio* species show an amazing variety of colour patterns, which range from horizontal stripes in *D. rerio* (Fig. 1a), over vertical bars in *D. aesculapii, D. choprae* or *D. erythromicron* (Fig. 1c, g, m) to spotted patterns in *D. tinwini* or *D. margaritatus* (Fig. 1d, h). The *Danio* species diversified for at least 13 million years in Southeast Asia and their spatial distributions only partially overlap today^10,45^. Hybrids between *D. rerio* and other *Danio* species can be produced in the laboratory by natural matings or by *in vitro* fertilization^19^. They invariably display colour patterns similar to the stripes in *D. rerio*, thus, horizontal stripes appear to be dominant over divergent patterns (Fig. 1e, f, i, j)^19^; whether this is due to a gain-of-function in striped species or losses in the other species is an open question^25,28,29^. The hybrids are virtually sterile impeding further genetic experiments, like QTL mapping, but they allow interspecific complementation tests^19^. Using this approach, we identified the potassium channel gene *Kcnj13* as repeatedly evolved in the *Danio* genus.

### Prevalent pattern orientation by the horizontal myoseptum

To reconstruct the history of colour pattern evolution we first investigated how pattern orientation is inherited in hybrids (see phylogenetic relationships depicted on the left of Fig. 4). The horizontal orientation of the stripes in *D. rerio* depends on the horizontal myoseptum (Fig. 1a, b)^11,13^. The closest sibling species to *D. rerio, D. aesculapii*, shows a very different pattern of vertically oriented dark bars (Fig. 1c)^8^. Similar barred patterns are exhibited by the more distantly related *D. choprae* and *D. erythromicron* (Fig. 1g, m). These patterns clearly do not use the horizontal myoseptum for orientation. In all three cases, hybrids with *D. rerio* show a pattern that resembles the horizontal *D. rerio* stripes (Fig. 1e, i, n)^29^. Strikingly, hybrids between *D. aesculapii* and *D. choprae* display a barred pattern (Fig. 1k). This indicates that the cues for horizontal orientation are lacking and that the pattern develops in a similar manner in both barred species. In contrast, hybrids between *D. aesculapii* and *D. erythromicron* develop highly variable patterns without any clear orientation (Fig. 1o; Extended Data Fig. 1). Therefore, the vertical bars must develop in a different manner in *D. erythromicron* compared to *D. aesculapii* and *D. choprae*.

Two *Danio* species display spotted patterns: *D. tinwini* has dark spots on a light background (Fig. 1d)^9^, whereas *D. margaritatus* shows light spots on a dark background (Fig. 1h)^7^. In both cases, hybrids with *D. rerio* show a stripe pattern similar to *D. rerio* (Fig. 1f, j)^29^. Hybrids between the two spotted species also develop a pattern of horizontal stripes, albeit with some interruptions and irregularities (Fig. 1l). These results indicate that the horizontal myoseptum functions to orient the pattern in the hybrids between *D. tinwini* and *D. margaritatus*, and therefore in at least one of the two parental species. It seems likely that this is the case in *D. tinwini*, as the spots show some horizontal orientation reminiscent of interrupted stripes. Hybrids between *D. aesculapii* and *D. margaritatus* develop meandering patterns that do not resemble either of the parental species and lack a clear horizontal or vertical orientation (Fig. 1p). Based on the most recent phylogeny^10^, we hypothesize an evolutionary history, in which the horizontal orientation of the pattern in the *D. rerio* group was gained from an ancestral ambiguous pattern and lost again in *D. aesculapii*. Two other species, *D. erythromicron* and *D. choprae*, independently might have acquired a vertical orientation from this ancestral pattern. The patterns of the hybrids between *D. aesculapii* and *D. erythromicron* or *D. margaritatus*, which are without clear orientation, might resemble such an ancestral pattern. These patterns are much more variable than the species patterns (Extended Data Fig. 1) suggesting that the ancestral patterns did not function as recognition signals but rather provided camouflage.

### Chromatophore interactions in stripes and bars

To investigate the developmental and genetic basis for the differences in pattern formation, we focussed on the sibling species *D. rerio* and *D. aesculapii*, which display horizontal stripes and vertical bars, respectively (Fig. 1a, c). In *D. rerio*, during early metamorphosis, iridophores emerge along the horizontal myoseptum to form the first light stripe (Extended Data Fig. 2a)^11-13^. In contrast, in *D. aesculapii* iridophores appear more scattered over the flank and only during later stages (Extended Data Fig. 2b, d). This indicates that it is not the physical presence of the horizontal myoseptum, which exists in both species, but specific guidance signals directing the cells into the skin in *D. rerio*, which lack in *D. aesculapii*. Later, when iridophores, covered by compact xanthophores, have formed the first contiguous light stripe with adjacent melanophore stripes in *D. rerio* (Extended Data Fig. 2c, e), in *D. aesculapii* melanophores and xanthophores intermix broadly (Extended Data Fig. 2f); they sort out loosely into vertical bars of low contrast without coherent sheets of dense iridophores between the melanophore bars during later stages (Extended Data Fig. 2h). Our observations suggest that the different patterns in these sibling species are produced by the presence or absence of guidance signals for iridophores along the horizontal myoseptum as well as by cellular interactions that prevent mixing of melanophores and xanthophores in *D. rerio* but not in *D. aesculapii*.

To address the role of the different cell types, we used the CRISPR/Cas9 system to generate mutants lacking individual chromatophore types in *D. aesculapii*. Whereas in *D. rerio* vestiges of the striped pattern form in the absence of one chromatophore type (Fig. 2a, b, c)^11^, loss of either melanophores (Fig. 2d) or xanthophores (Fig. 2e) completely abolishes the patterning in *D. aesculapii*. This indicates that the repulsive interactions between melanophores or xanthophores and iridophores, which account for the residual patterns in *D. rerio*^11,12^, are absent in *D. aesculapii*. In contrast, eliminating iridophores in *D. aesculapii* still permits some melanophore bar formation (Fig. 2f). This indicates that iridophores, which play a dominant role for stripe formation in *D. rerio*, are dispensable for the formation of vertical bars in *D. aesculapii*.

**Fig. 2:**
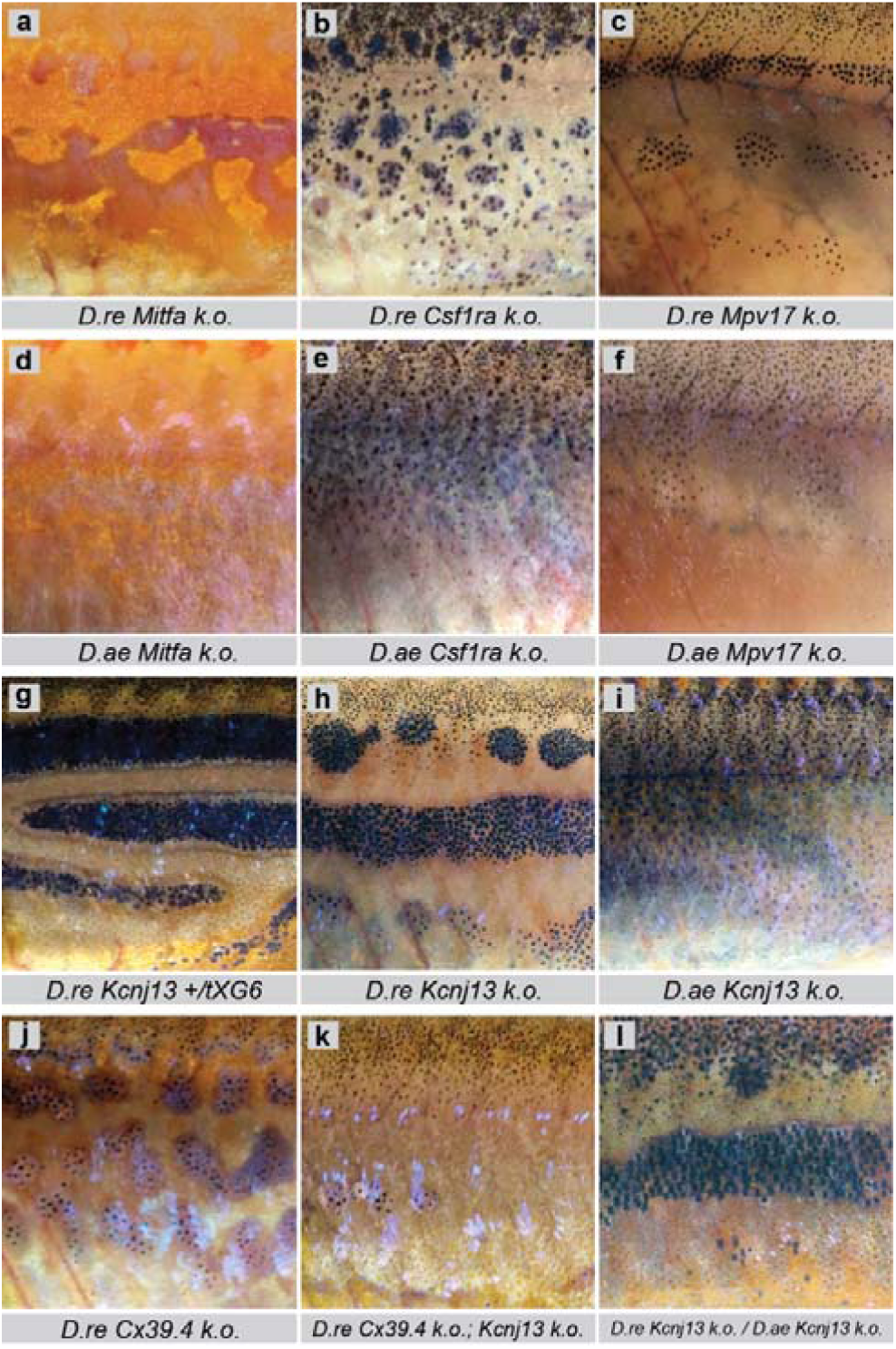
Mutant phenotypes in *D. rerio, D. aesculapii* and their hybrids. In *D. rerio* loss of one type of pigment cell, **a**, melanophores in *Mitfa* (*nacre*) mutants, **b**, xanthophores in *Csf1ra* (*pfeffer*) mutants, or **c**, iridophores in *Mpv17* (*transparent*) mutants, still permits rudimentary aggregation of dense iridophores (a) or melanophores (b, c). In *D. aesculapii*, **d**, loss of melanophores in *Mitfa* mutants or **e**, loss of xanthophores in *Csf1ra* mutants, abrogate any residual pattern formation. However, **f**, bars still form in *Mpv17* mutants, despite the absence of iridophores. **g**, *D. rerio* heterozygous for the dominant allele *Kcnj13*^*tXG6*^*/obelix*. **h**, *D. rerio* homozygous for the loss-of-function allele *Kcnj13*^*t24ui*^. **i**, *D. aesculapii* homozygous for a *Kcnj13* loss-of-function allele. **j**, *D. rerio* homozygous for the loss-of-function of Cx39.4. **k**, in *D. rerio* double mutants for *Cx39*.*4* and *Kcnj13* patterning is completely absent. **l**, interspecific hybrids between *D. rerio* and *D. aesculapii* that are both mutant in *Kcnj13*.

Next, we analysed genes with known functions in heterotypic interactions between chromatophores. In *D. rerio* loss-of-function mutations in the gap junction genes *Cx39*.*4* (*luchs*)^44,46^ and *Cx41*.*8/Gja5b* (*leopard*)^41,46,47^ as well as mutations in *Igsf11* (*seurat*)^42^, which codes for a cell adhesion molecule, lead to melanophore spots (Fig. 2j, Extended Data Fig. 3a, c, e), whereas mutations in *Kir7*.*1/Kcnj13* (*obelix*/*jaguar*) result in fewer and wider stripes with some mixing of melanophores and xanthophores (Fig. 2g)^20,48^. So far, only dominant alleles of *Kcnj13* have been described^41,46,49^. We used the CRISPR/Cas9 system to generate a *Kcnj13* loss-of-function allele in *D. rerio*, which is recessive (Fig. 2h, Extended Data Fig. 3e, g, Supplementary Information). To investigate the functions of these four genes in *D. aesculapii*, we generated mutants in the orthologs. In all of them we find an even distribution of melanophores (Fig. 2i, Extended Data Fig. 3b, d, f, h) indicating that the interactions mediated by these genes are essential to generate the melanophore bars in *D. aesculapii*. The complete loss of a pattern in single mutants in *D. aesculapii* is different from *D. rerio* where this occurs only in double mutants (Fig. 2k)^46^. In concert with predictions of agent-based models of patterning^50^, this indicates that the robust formation of horizontal stripes in *D. rerio* is due to a gain in complexity based on partially redundant chromatophore interactions. These are dominated by iridophores and oriented by an as yet unidentified signal along the horizontal myoseptum. *D. aesculapii* might have secondarily lost the dominance of iridophores leading to a pattern based primarily on interactions between xanthophores and melanophores and thus of lower complexity.

The ability to generate loss-of-function mutations in both species allowed us to generate interspecific hybrids, which carry loss-of-function alleles from both parental species. These hybrids are very similar to the respective *D. rerio* mutants (Fig. 2l, Extended Data Fig. 3i, j), indicating that these genes have the same functions during stripe formation in *D. rerio* and the hybrids.

### *Kcnj13* evolved repeatedly in the *Danio* genus

Next, we generated reciprocal heterozygotes, i.e. interspecific hybrids carrying a loss-of-function allele from either one of the parental species in an otherwise identical genetic background^18^. This powerful genetic test to identify evolved genes has not been applied in vertebrates, so far. We expect similar patterns in these hybrids if the gene function can be fully provided by either wild-type allele. A qualitatively altered hybrid pattern would reveal that one of the wild-type alleles cannot complement the loss-of-function of the other, therefore indicating functional changes during evolution. We found that heterozygous hybrids with the loss-of-function allele of *Kcnj13* from *D. rerio* display a pattern of spots or interrupted stripes whereas a striped pattern forms with the mutant allele from *D. aesculapii* (Fig. 3a, b, Extended Data Fig. 4g, h). This indicates that the wild-type allele from *D. aesculapii* cannot compensate for the loss of the *D. rerio* allele. In contrast, in the case of heterozygous hybrids with *Cx39*.*4, Cx41*.*8* and *Igsf11* striped patterns indistinguishable from wild-type hybrids are formed regardless whether the wild-type allele stems from *D. rerio* (Extended Data Fig. 4b, d, f) or the other species (Extended Data Fig. 4a, c, e). These reciprocal heterozygosity tests indicate that *Cx39*.*4, Cx41*.*8* and *Igsf11* provide similar functions in both species, whereas the function of *Kcnj13* has evolved between the two species.

**Fig. 3:**
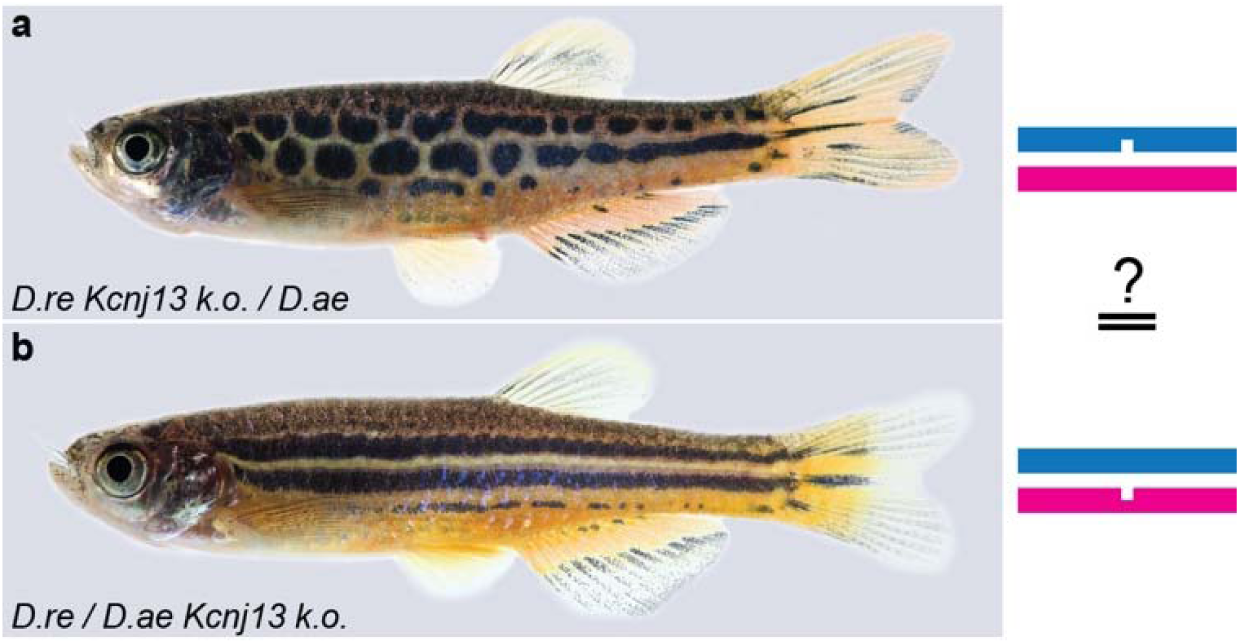
A reciprocal heterozygosity test to identify *Kcnj13* evolution. Two interspecific hybrids between *D. rerio* and *D. aesculapii*, which are heterozygous for a *Kcnj13* loss-of-function mutation. **a**, stripes are interrupted in hybrids carrying the mutant allele from *D. rerio* (nick in the blue line representing the zebrafish genome). **b**, hybrids carrying the mutant allele from *D. aesculapii* (nick in the magenta line, representing the *D. aesculapii* genome) are indistinguishable from wild-type hybrids (Fig. 1e).

To investigate if *Kcnj13* underlies the pattern variation more broadly across the *Danio* genus, we tested seven additional species (Fig. 4a, top to bottom). As mentioned above, wild-type hybrids between *D. rerio* and all other *Danio* species display horizontal stripes, resembling the *D. rerio* pattern, with slight defects in *D. albolineatus* (Fig. 4b). Strikingly, not only *D. rerio Kcnj13* k.o. / *D. aesculapii* hybrids (Fig. 4c, highlighted in magenta) but also *D. rerio Kcnj13* k.o. / *D. tinwini* hybrids developed patterns of spots or interrupted stripes indicating that the *Kcnj13* function must have evolved compared to *D. rerio* (Fig. 4c, highlighted in yellow). This pattern, which is qualitatively different from all wild-type hybrids and also from the *D. rerio Kcnj13* mutant pattern, is similar to the parental pattern of *D. tinwini*, where dense iridophores interrupt the dark melanophore stripes (Fig. 1d, Fig. 4c). *D. rerio Kcnj13* k.o. / *D. choprae* hybrids also developed patterns that resemble interrupted stripes (Fig. 4c, highlighted in cyan), similar to the *D. rerio Kcnj13* k.o. / *D. aesculapii* hybrid pattern (Fig. 4c). No qualitative differences were detected between wild-type hybrids (Fig. 4b) and hybrids heterozygous for *D. rerio Kcnj13* in the case of *D. kyathit, D. nigrofasciatus, D. albolineatus, D. erythromicron* and *D. margaritatus* (Fig. 4c). This indicates that the alleles from these species complement the loss of the *D. rerio Kcnj13* allele. Functional changes of *Kcnj13* occurred in *D. aesculapii, D. tinwini* and *D. choprae* compared to *D. rerio*, however, heterozygous hybrids did not develop pure *D. rerio Kcnj13* mutant patterns indicating that the orthologs still provide some function for patterning across all species tested. The separated positions of the three species with the different functions of *Kcnj13* in the phylogenetic tree (graph on the left of Fig. 4)^10^ indicate a repeated and independent evolution of the same gene.

**Fig. 4:**
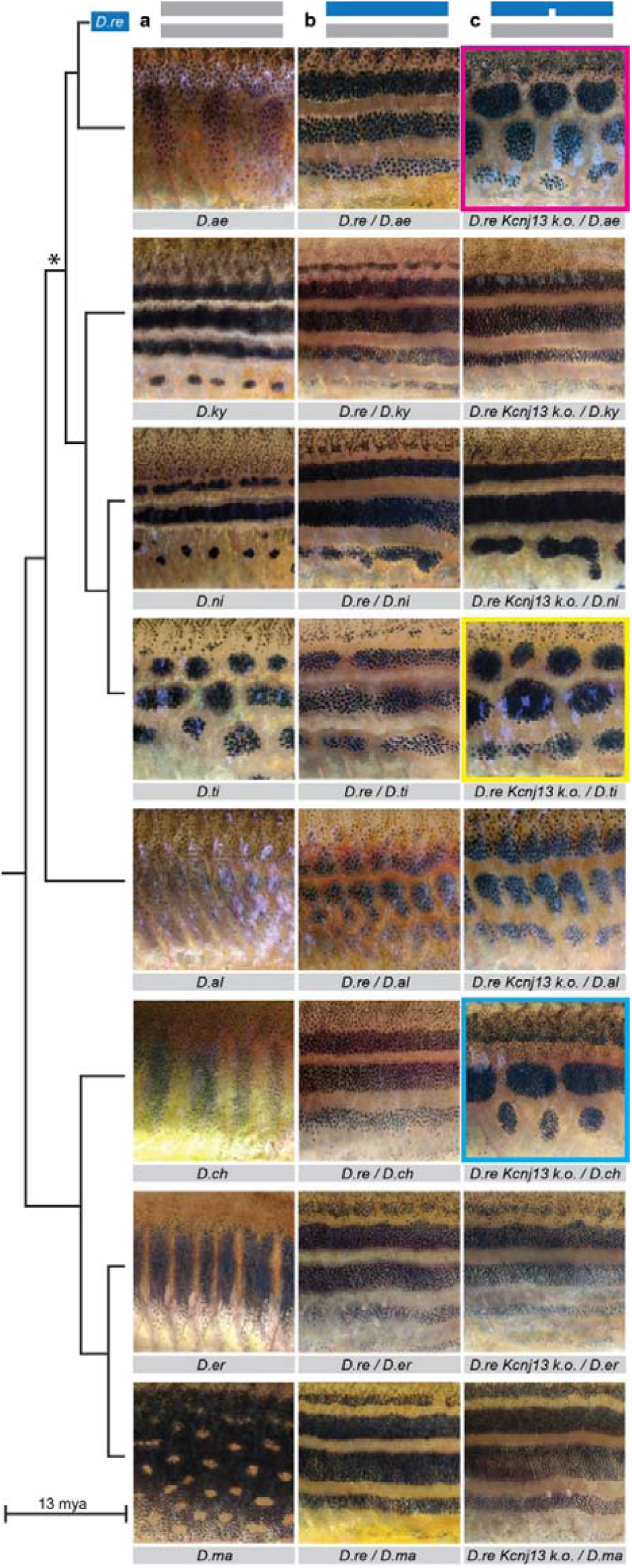
Repeated *Kcnj13* evolution. **a**, From top to bottom wild-type *Danio* colour patterns (*D. rerio* species group: barred *D. aesculapii, D*.*ae*, the closest relative to *D. rerio*; striped *D. kyathit, D*.*ky*; striped *D. nigrofasciatus, D*.*ni*; spotted *D. tinwini, D*.*ti*;) unpatterned *D. albolineatus, D*.*al*; (*D. choprae* species group: barred *D. choprae, D*.*ch*; barred *D. erythromicron, D*.*er*; spotted *D. margaritatus, D*.*ma*). The phylogenetic tree to the left depicts the relationships between the species^1^. The asterisk denotes an uncertainty in the phylogenetic relationships within the *D. rerio* species group. **b**, Hybrids between *D. rerio* (*D*.*re*) and eight other danios (from top to bottom *D. aesculapii, D*.*ae, D. kyathit, D*.*ky, D. nigrofasciatus, D*.*ni, D. tinwini, D*.*ti, D. albolineatus, D*.*al, D. choprae, D*.*ch, D. erythromicron, D*.*er, D. margaritatus, D*.*ma*). **c**, Interspecific hybrids carrying the *Kcnj13* k.o. allele from *D. rerio* and the wild-type allele in the eight species. Pattern defects occur in three cases, *D. rerio Kcnj13* k.o. */ D. aesculapii* (magenta box), *D. rerio Kcnj13* k.o. */ D. tinwini* (yellow box) and *D. rerio Kcnj13* k.o. */ D. choprae* (cyan box). In the other five cases the patterns in heterozygous hybrids do not differ from the striped patterns of wild-type hybrids.

### Kcnj13

Potassium channels have important roles in tissue patterning^51^, notably in the regulation of allometric growth of fins in *D. rerio*^52,53^. *Kcnj13* encodes an inwardly rectifying potassium channel (Kir7.1) conserved in vertebrates (Extended Data Fig. 5). Mutations are known to cause defects in tracheal development in mice^54^ and two rare diseases in humans leading to visual impairment^54-60^. During colour pattern formation in *D. rerio* its function is autonomously required in melanophores^48^, and in *ex vivo* studies it was shown that the channel is involved in the contact-dependent depolarisation of melanophores upon interaction with xanthophores leading to a repulsion between these cells^43^. Evolution in *Kcnj13* in *D. aesculapii, D. tinwini* and *D. choprae* might therefore cause differences in heterotypic chromatophore interactions between species. The protein coding sequences of *Kcnj13* orthologs are highly conserved in all *Danio* species with only very few diverged sites in the cytoplasmic N- and C-terminal parts of the protein (Extended Data Fig. 6). Whether any of these amino acid changes might affect the function of the channel and/or if changes in gene expression are the basis for the repeated evolution of *Kcnj13* will require further experiments.

In contrast to mammals and birds, basal vertebrates retained several chromatophore types providing a substrate for the development of elaborate colour patterns. Their rapid and extensive evolutionary diversification is most likely if the number of underlying genes is small^61^. Several patterning genes have been repeatedly identified in genetic screens in *D. rerio*^41,46,49^. These genes provide candidates that might have evolved to contribute to patterning differences between *Danio* species. Evolved genes have been identified in *D. albolineatus* and *D. nigrofasciatus*, where changes in two signalling pathways, Csf1 or Edn3, likely underlie patterning variations by differentially promoting xanthophore or iridophore development, respectively^19,36,62^. We focused on genes regulating heterotypic interactions between chromatophores as a potential genetic basis for colour pattern evolution. Using interspecific mutant complementation tests we identified the potassium channel gene *Kcnj13* as contributing to patterning divergence in multiple *Danio* species. We have shown that this genus offers the opportunity to identify evolved genes and to reconstruct the evolution of biodiversity.

## Methods

No statistical methods were used to predetermine sample size. The experiments were not randomized. The investigators were not blinded to allocation during experiments and outcome assessment.

### Fish husbandry

Zebrafish, *D. rerio*, were maintained as described earlier^1^. If not newly generated (Table 4, Supplementary Information), the following genotypes were used: wild-type Tuebingen/TU, *nacre*^*w2*^*/nac/Mitfa*^2^, *pfeffer*^*tm236*^*/pfe/Csf1ra*^3^, *transparent*^*b6*^*/tra/Mpv17*^4^, *leopard*^*t1*^*/leo/Cx41*.*8*^5,6^, *luchs* ^*t37ui*^*/luc/Cx39*.4^7^, *obelix*^*tXG6*^*/obe/Kcnj13*^7^.

*D. aesculapii* and *D. albolineatus* were treated identical to *D. rerio*. For the other *Danio* species, *D. kyathit, D. tinwini, D. nigrofasciatus, D. choprae, D. margaritatus* and *D. erythromicron* individual pair matings were not successful. Therefore, the fish were kept in groups in tanks containing boxes lightly covered with Java moss (*Taxiphyllum barbieri*), which resulted in sporadic matings and allowed us to collect fertilized eggs.

Interspecific hybrids were either obtained by natural matings or by in vitro fertilizations^8^. Heterozygous or trans-heterozygous mutant hybrids were identified by PCR and sequence analysis using specific primer pairs (Table 1 and 3, Supplementary Information).

All species were staged according to the normal table of *D. rerio* development^9^. All animal experiments were performed in accordance with the rules of the State of Baden-Württemberg, Germany, and approved by the Regierungspräsidium Tübingen.

### CRISPR/Cas9 gene editing

The CRISPR/Cas9 system was applied either as described in^10^ or according to the guidelines for embryo microinjection of IDT. Briefly, oligonucleotides were cloned into pDR274 to generate the sgRNA vector (Table 2, Supplementary Information). sgRNAs were transcribed from the linearized vector using the MEGAscript T7 Transcription Kit (Invitrogen). Alternatively, target-specific crRNAs and universal tracrRNAs were purchased from IDT. sgRNAs or crRNA:tracrRNA duplexes were injected as ribonucleoprotein complexes with Cas9 proteins into one-cell stage embryos. The efficiency of indel generation was tested on eight larvae at 1 dpf by PCR using specific primer pairs and by sequence analysis as described previously (Table 1 and 3, Supplementary Information)^11^. The remaining larvae were raised to adulthood. Mature F0 fish carrying indels were outcrossed. Loss-of-function alleles in heterozygous F1 fish were selected to establish homozygous or trans-heterozygous mutant lines (Table 4, Supplementary Information).

### Image acquisition

Anesthesia of adult fish was performed as described previously^11^. A Canon 5D Mk II camera was used to obtain images. Juvenile fish were either embedded in low melting point agarose or fixed in 4% formaldehyde/0.08% glutaraldehyde and then photographed under a Leica MZ1 stereomicroscope (Extended Data Fig. 2). Images were processed using Fiji^12^, Adobe Photoshop and Adobe Illustrator CS6.

### RNA-Sequencing and transcriptome analysis

#### Skin of *Danio* species

Adult fish (n=5 each for *D. rerio* (TU), *D. aesculapii, D. kyathit, D. nigrofasciatus, D. tinwini, D. albolineatus, D. choprae, D. erythromicron, D. margaritatus*) were euthanized by exposure to buffered 0.5 g/L MS-222 (Tricaine). Skin tissues were dissected in ice-cold PBS and collected using TRIzol (Life Technologies). RNA integrity and quantity were assessed by Agilent 2100 Bioanalyzer. Library preparation (TruSeq stranded mRNA, Illumina; 200 ng per sample) and sequencing (NovaSeq 6000, 2 × 100 bp) were performed by CeGaT GmbH (Tübingen, Germany). RNA-Seq analysis was carried out using the *Danio rerio* GRCz11 genome build for all *Danio* species and STAR aligner with default settings^13^. Differential expression analysis was then carried out using DESeq2^14^. The actual commands used can be found here: https://github.com/najasplus/STAR-deseq2.

### Sequence analysis

We found SNPs in the coding region of *Kcnj13* and considered other resources^15^, including the latest zebrafish reference genome assembly (GRCz11) and the ENA deposition Zebrafish Genome Diversity (PRJEB20043, Wellcome Trust Sanger). Also, all identified SNPs in the *kcnj13* coding sequence from the Zebrafish Mutation Project were incorporated^16^. The variant calling pipeline for all *Danio* species consisted of GATK 3.8 and 4 and picard^17^ from STAR-aligned bam files based on GATK Best-Practices pipeline. The full commands used can be found here: https://github.com/najasplus/rnaseq_variant_calling. Furthermore, variants were also called and checked using SAMtools, mpileup and bcftools^18^. The protein sequence alignment was produced using T-coffee^19^ and refined using BOXSHADE (developed by Kay Hofmann and Michael D. Baron, unpublished) and Microsoft Word.

## Data and code availability

The dataset generated during this study is available at The European Nucleotide Archive (ENA) accession number: pending.

## Acknowledgements

We thank all members of the Nüsslein-Volhard laboratory. This work was supported by an ERC Advanced Grant “DanioPattern” (694289) and the Max Planck Society, Germany.

## Contributions

All authors were involved in the design of the experiments. M.P., U.I. and H.G.F. performed the experiments. U.I., C.N.V., M.P., H.G.F. and C.D. analysed the data with support of A.E. M.P. made the figures with contributions from U.I. and C.N.V. U.I., C.N.V. and M.P. wrote the manuscript. C.N.V. and U.I. acquired funding.

## Ethics declaration

Competing interests

The authors declare no competing interests.

## Extended Data Figures, Tables and Legends

**Extended Data Figure 1:**
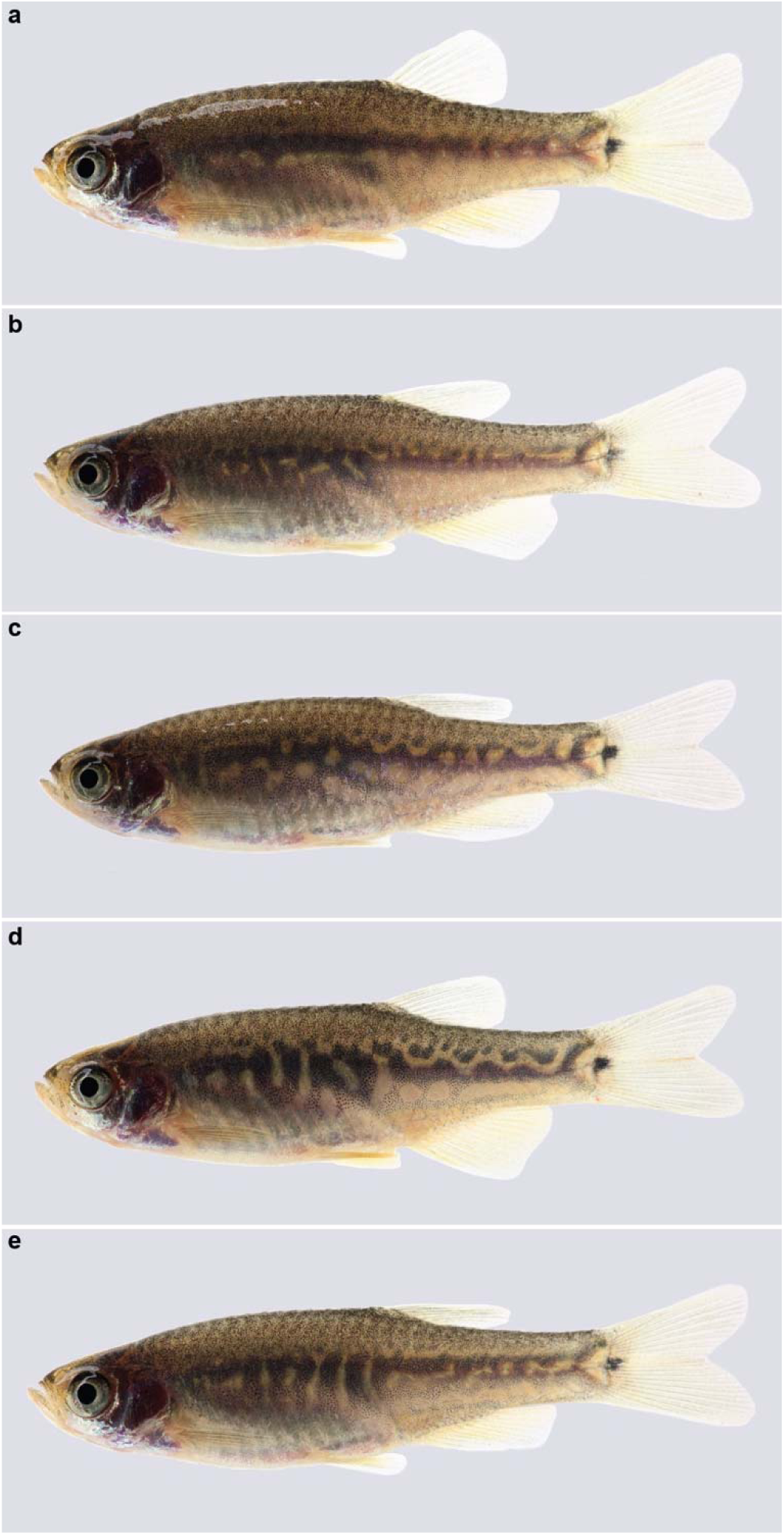
Variability of patterns in interspecific hybrids between *D. aesculapii* and *D. erythromicron*. Interspecific hybrids between *D. aesculapii* and *D. erythromicron*, **a - e**, show a range of patterns without clear horizontal or vertical orientation.

**Extended Data Figure 2:**
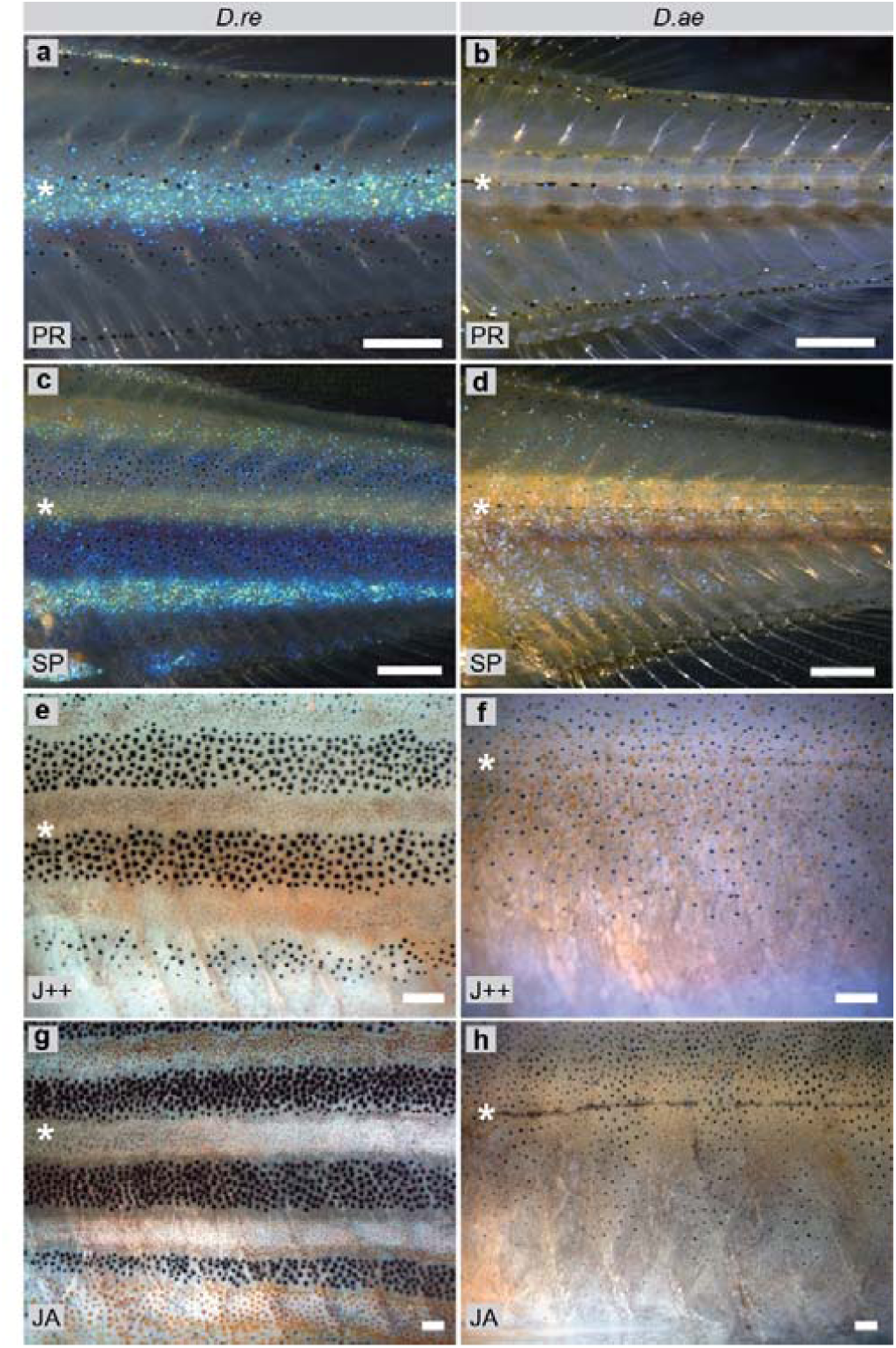
Development of colour patterns in *D. rerio* and *D. aesculapii*. **a**, *D. rerio* fish at stage PR. Iridophores (arrowhead) emerge along the horizontal myoseptum (asterisk) to form the first light stripe. **b**, *D aesculapii* fish at stage PR. **c**, *D. rerio* at stage SP. The first light stripe is flanked dorsally and ventrally by dark stripes **d**, *D. aesculapii* at stage SP. Iridophores emerge in a scattered fashion. **e**, *D. rerio* at stage J++. Light stripes are covered by compact xanthophores **f**, *D. aesculapii* at stage J++. Melanophores and xanthophores broadly intermix. **g**, *D. rerio* at stage JA. **h**, *D. aesculapii* at stage JA. Melanophores and xanthophores sort out loosely into vertical bars of low contrast; no dense iridophores are visible between the dark bars. a-d: incident light illumination to highlight iridophores, e-h: bright field illumination to visualise xanthophores and melanophores. Staging according to^2^. PB (pectoral fin bud, 7.2 mm SL). SP (squamation posterior, 9.5 mm SL). J++ (juvenile posterior, 16 mm SL). JA (juvenile-adult, >16 mm SL). Scale bars correspond to 250 μm.

**Extended Data Figure 3:**
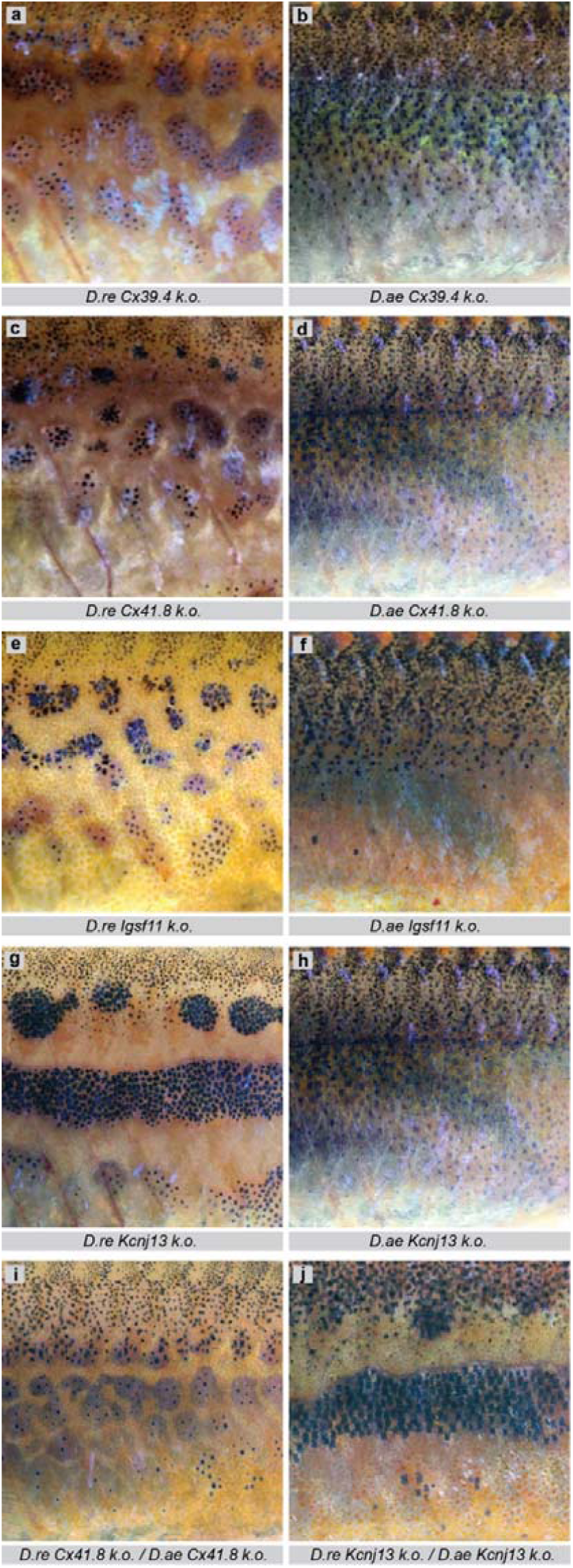
Mutant phenotypes in *D. rerio* and *D. aesculapii* of genes required for heterotypic interactions. **a**, *D. rerio* (*D*.*re*) *Cx39*.*4* knock-out (k.o.) mutant^3^. **b**, *D. aesculapii* (*D*.*ae*) *Cx39*.*4* k.o. mutant. **c**, *D*.*re Cx41*.*8* mutant^4,5^. **d**, *D*.*ae Cx41*.*8* k.o. mutant. **e**, *D*.*re Igsf11* k.o. mutant^6^. **f**, *D. ae Igsf11* mutant. **g**, *D. rerio Kcnj13* k.o. mutant^3,4,7^. **h**, *D*.*ae Kcnj13* k.o. mutant. **i**, Interspecific hybrids between *D*.*re Cx41*.*8* and *D*.*ae Cx41*.*8* mutants. **j**, Interspecific hybrids between *D*.*re Kcnj13* and *D*.*ae Kcnj13* mutants. In all cases, loss-of function (k.o.) alleles were created using CRISPR/Cas9 gene editing.

**Extended Data Figure 4:**
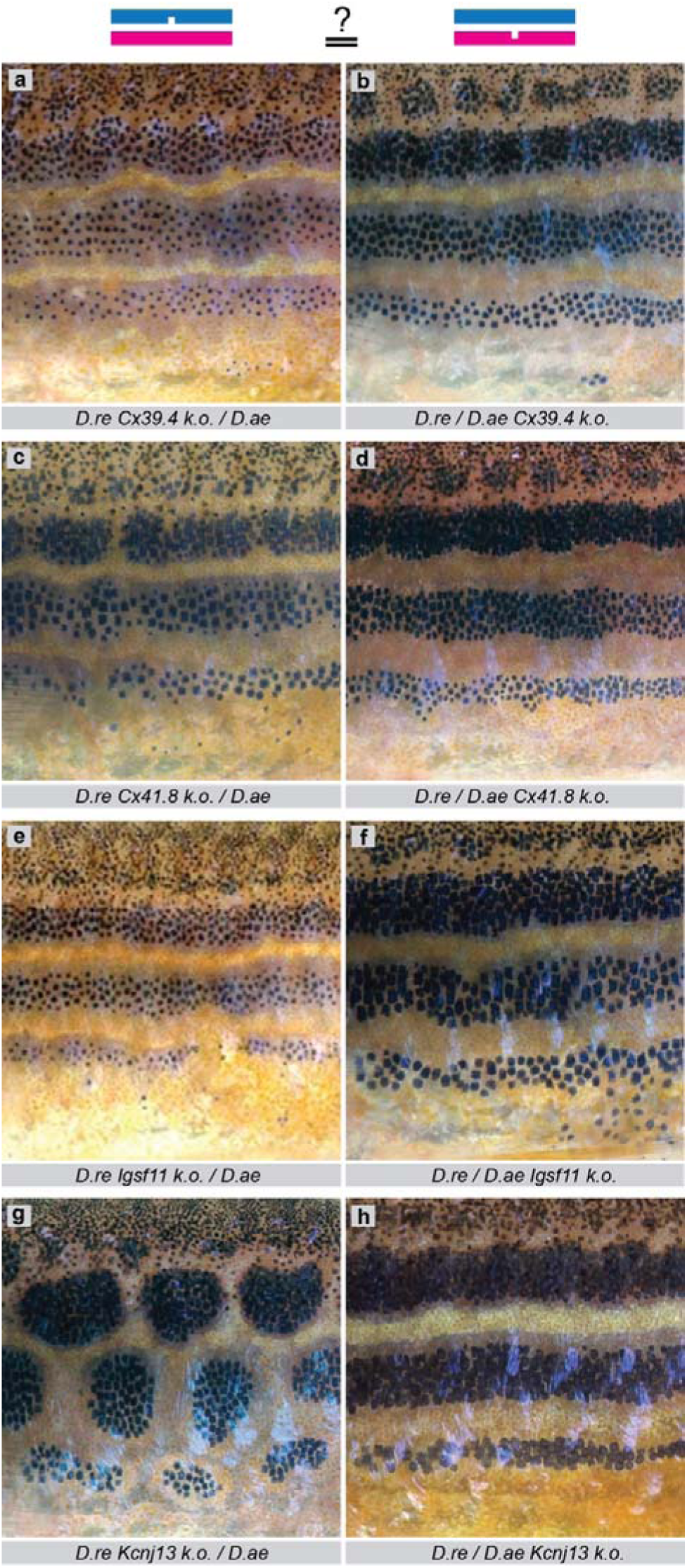
Reciprocal heterozygosity tests identify *Kcnj13* evolution. Interspecific heterozygous hybrids carrying a mutant allele (nicked bar) from either parental species (blue: *D. rerio, D*.*re*, or magenta: *D. aesculapii, D*.*ae*) in an otherwise identical genetic background. In the cases of **a**/**b**, *Cx39*.*4*, **c**/**d**, *Cx41*.*8* and **e**/**f**, *Igsf11* both hybrids show identical phenotypes. In the case of **g**/**h**, *Kcnj13*, the two hybrids show different phenotypes: Interrupted stripes and spots in those carrying the mutant allele from *D. rerio* (g) and a striped pattern in those carrying the mutant allele from *D. aesculapii* (h).

**Extended Data Figure 5:**
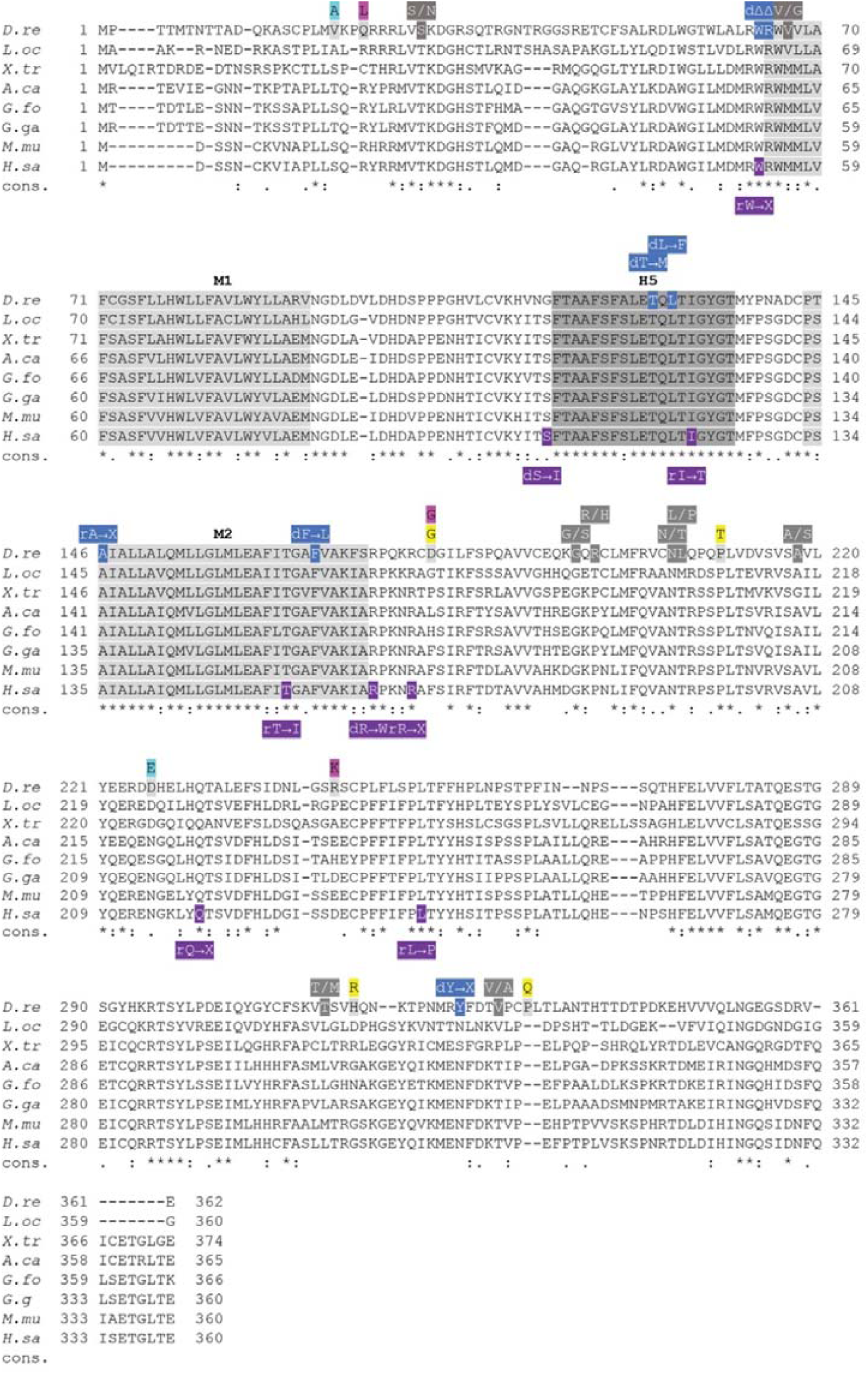
Kcnj13 sequence alignment of vertebrate orthologues. Sequence alignment of Kcnj13 orthologues from different vertebrate species. The two transmembrane domains (M1/M2) are shaded in light grey, the P-Loop (H5) in dark grey. Dominant (d) or recessive (r) mutations are indicated for zebrafish^3,4,7,8^ (blue) and human^9-13^ (purple). Positions that are different between *D. rerio* and *D. aesculapii* (magenta), *D. tinwini* (yellow), *D. choprae* (cyan) and polymorphic positions in *D. rerio* (dark grey) are highlighted. Kcnj13 sequences of *Danio rerio* (zebrafish, NP_001039014.1), *Lepisosteus oculatus* (spotted gar, XP_006638004.1), *Xenopus tropicalis* (tropical clawed frog, NP_001096437.1), *Anolis carolinensis* (green anole, XP_016847621.1), *Geospiza fortis* (medium ground finch, XP_005430275.1), *Gallus gallus* (chicken, XP_015132697.1), *Mus musculus* (house mouse, NP_001103697.1) and *Homo sapiens* (human, NP_002233.2).

**Extended Data Figure 6:**
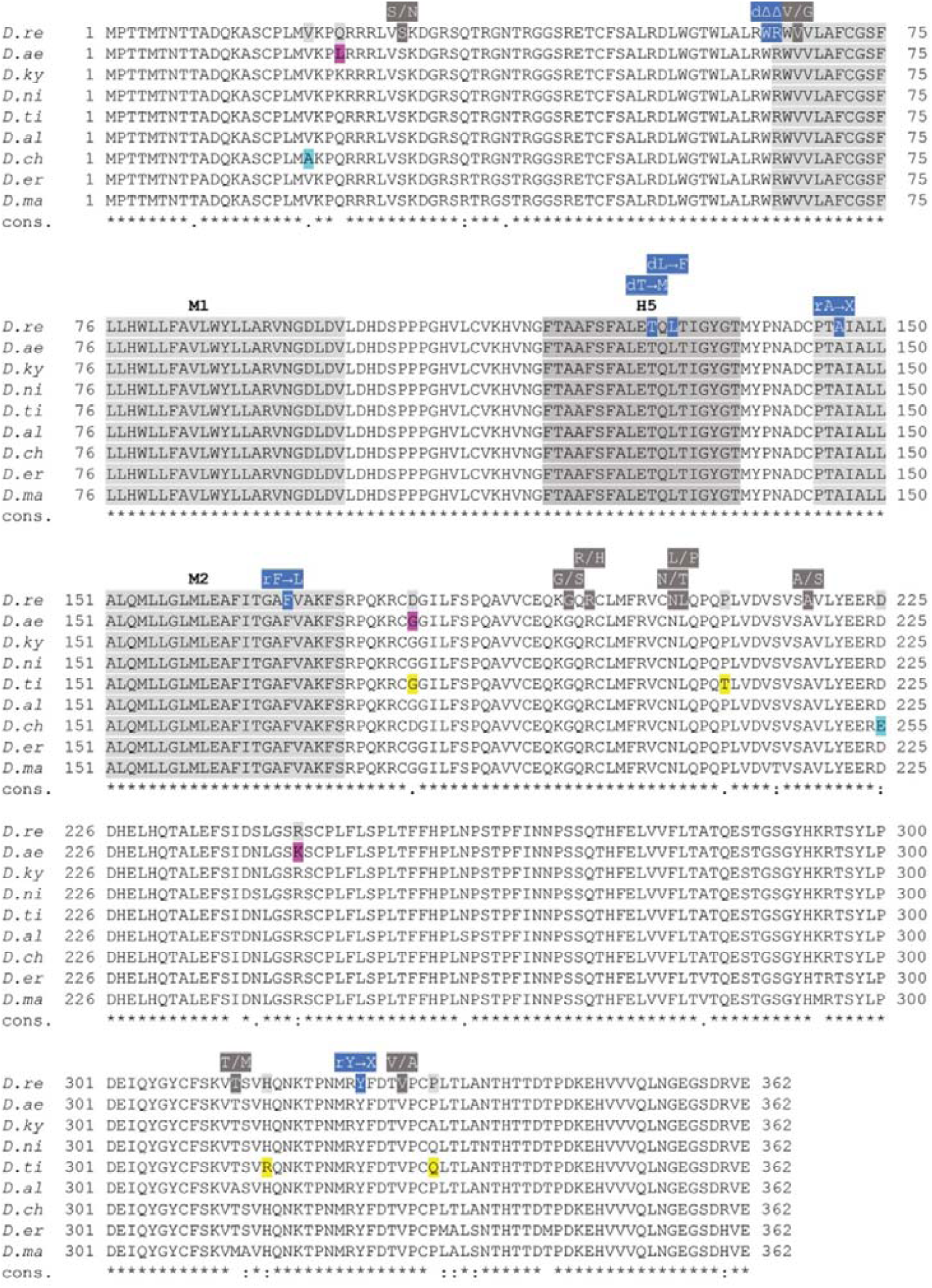
Sequence alignment of Kcnj13 orthologues from *Danio* species. Kcnj13 sequences from *D. rerio* (*D*.*re*), *D. aesculapii* (*D*.*ae*), *D. kyathit* (*D*.*ky*), *D. nigrofasciatus* (*D*.*ni*), *D. tinwini* (*D*.*ti*), *D. albolineatus* (*D*.*al*), *D. choprae* (*D*.*ch*), *D. margaritatus* (*D*.*ma*), *D. erythromicron* (*D*.*er*). Amino acids evolved between *D*.*re* and *D*.*ae* (magenta), *D*.*ti* (yellow) and *D*.*ch* (cyan). Dominant (d) or recessive (r) mutations in *D*.*re Kcnj13*^3,4,7,8^ (blue). Amino acid polymorphisms in *D*.*re* (dark grey). Transmembrane domains (M1/M2) (light grey blocks) and the P-loop (H5) (dark grey block).

## Supplementary Information and Legends

### Supplementary Tables

**Table 1.**
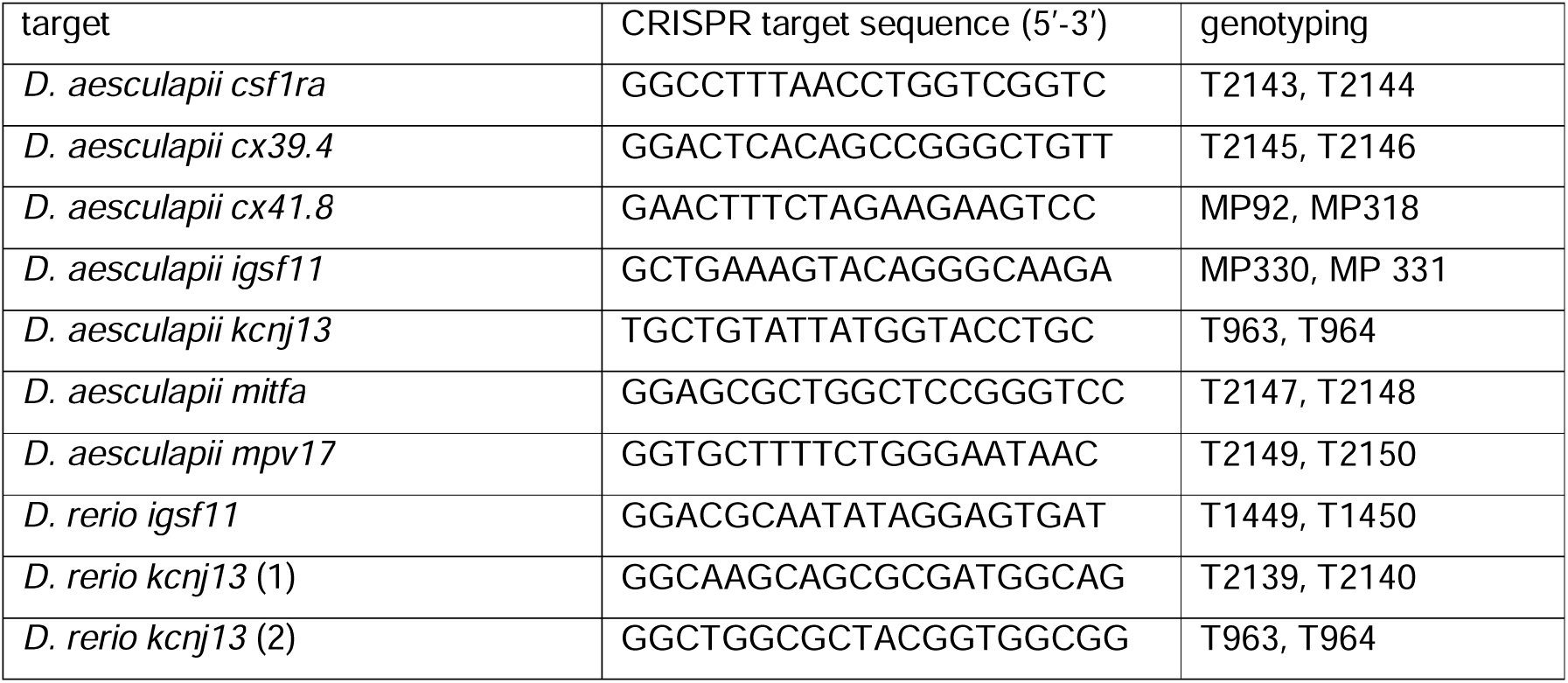
List of targeted genes.

**Table 2.**
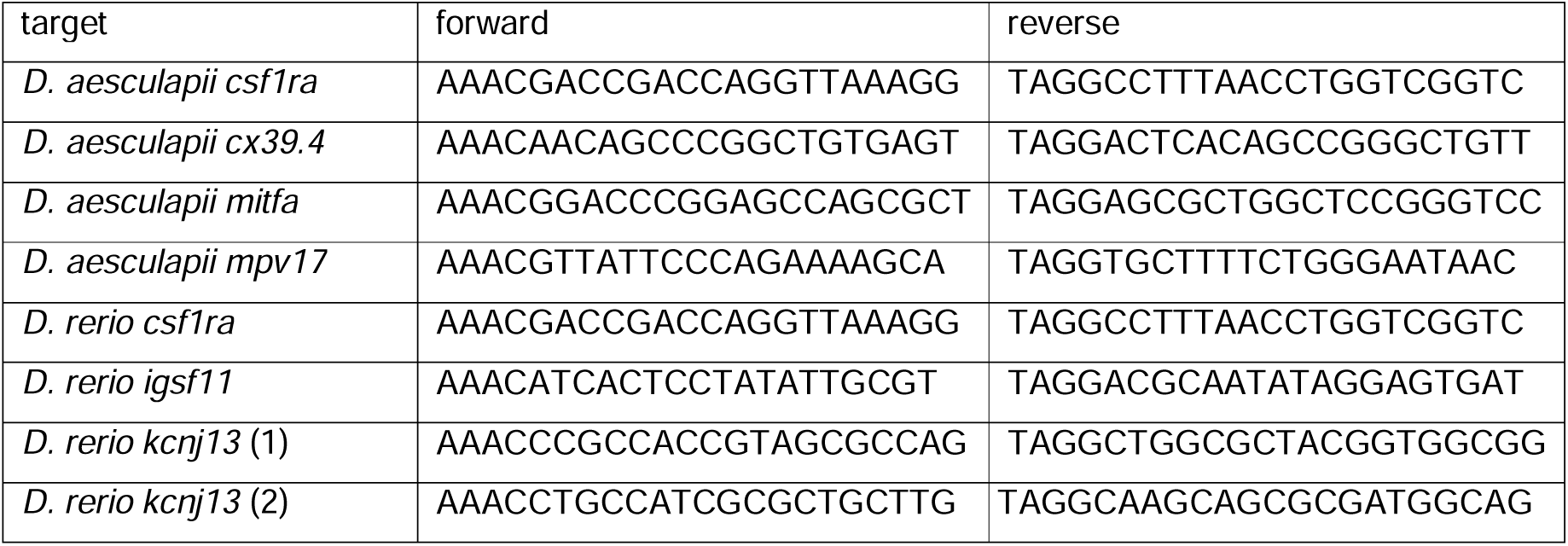
Primer pairs used for the generation of sgRNAs.

**Table 3.**
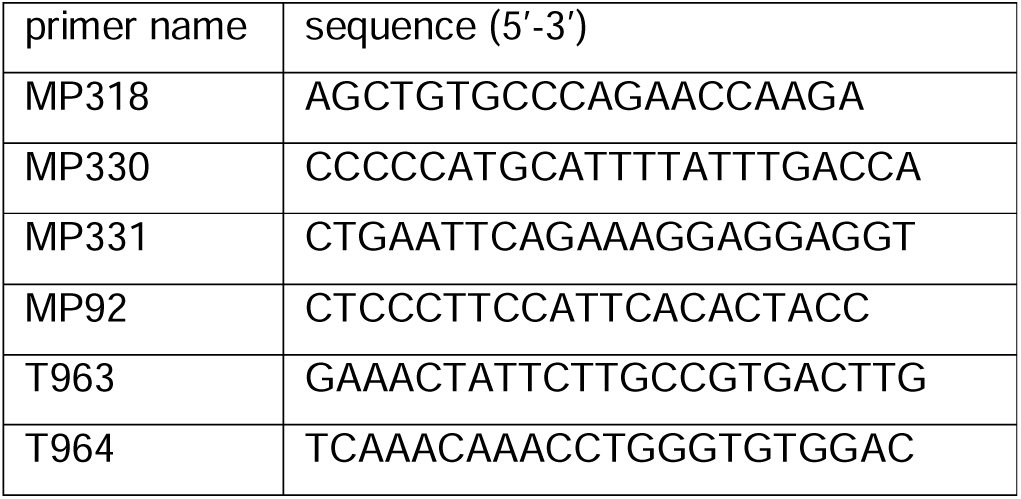
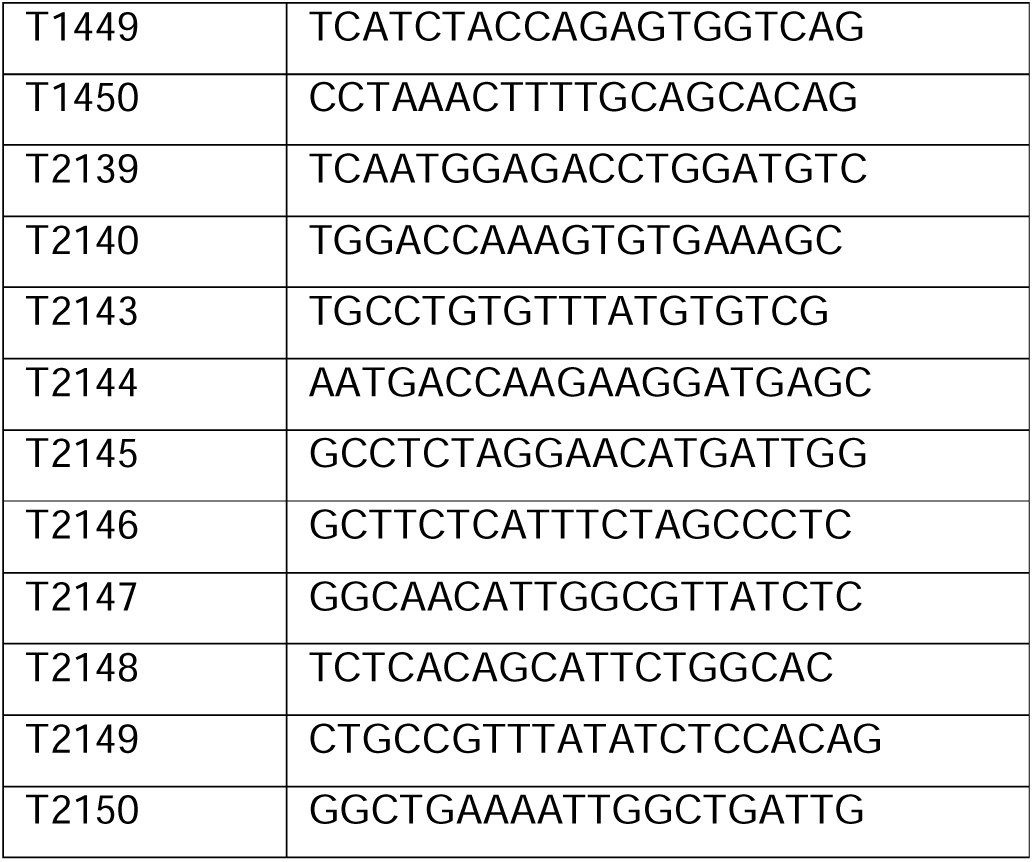
Primers used for genotyping.

**Table 4.**
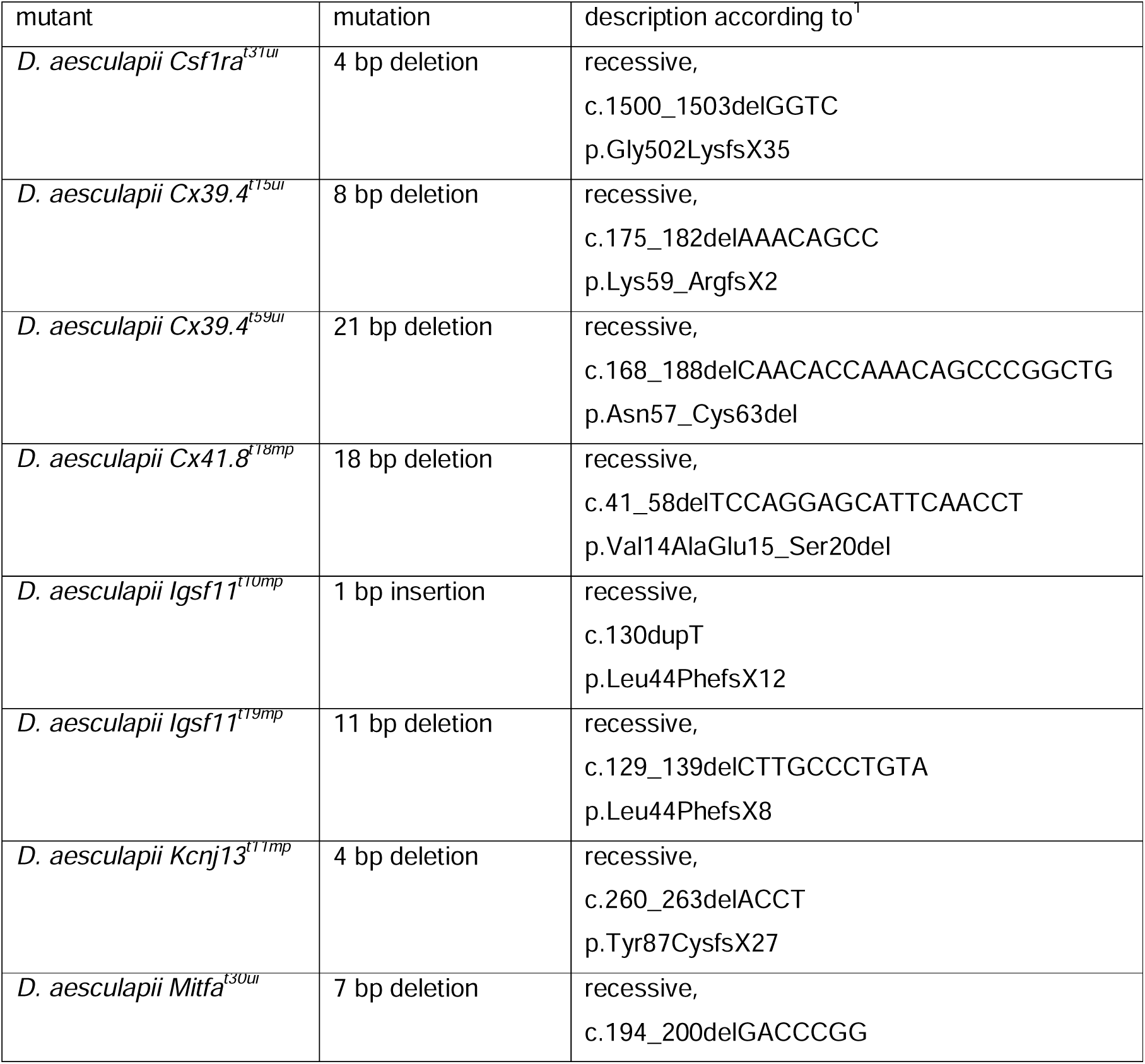
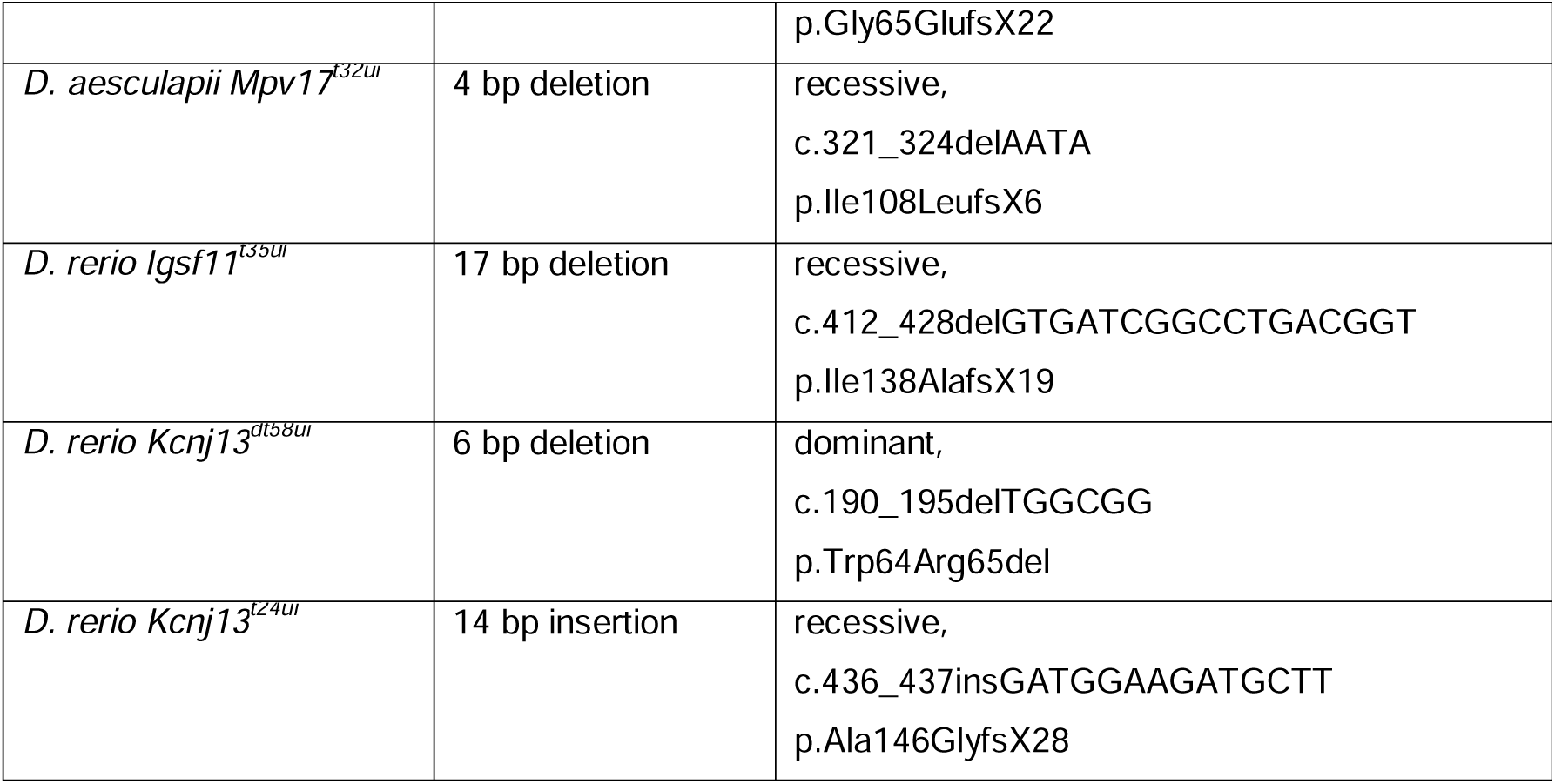
List of generated mutants.

### Supplementary Discussion

Kcnj13 functions as a tetramer (Fig. 4d, e), where each subunit contributes two transmembrane helices (M1 and M2, Extended Data Fig. 4) to the formation of the channel pore and a short extracellular loop that folds back to form the pore lining ion selectivity filter (P-loop or H5, Extended Data Fig. 4). The N- and C-termini of the subunits reside in the cytoplasm, where they also contribute to the ion pore, but are mainly involved in gating of the channel (reviewed in^2^). Four dominant *Kcnj13* alleles have been identified in *D. rerio* in several independent forward genetic screens (Extended Data Fig. 4)^3-5^. All of them show broad stripes with irregular interruptions when heterozygous (Fig. 2g) and strong pattern aberrations with fewer, wider and interrupted dark stripes and some mixing of melanophores and xanthophores when homozygous or trans-heterozygous. Three of them carry point mutations affecting H5 or M2^3,4,6^, one is the result of a C-terminal truncation (Extended Data Fig. 4)^5^. The point mutations lead to proteins that do not produce functional channels^6^ and it has been suggested that the dominant phenotype is caused by a dosage-dependent effect, i.e. haploinsufficiency^7^. To test this possibility we used the CRISPR/Cas9 system and generated a loss of function allele of *Kcnj13* in *D. rerio* which is recessive. The 14 base pair insertion near the end of the first coding exon leads to an early truncation of the protein. Homozygous mutants show a phenotype similar to homozygous mutants for the dominant alleles (Fig. 2h)^3-7^. This shows that the dominant alleles are dominant-negatives, where the mutant proteins inhibit the function of the wild-type protein in heterozygotes. The generation of this recessive allele allowed us to perform the reciprocal heterozygosity test with *D. aesculapii* as well as interspecific complementation with seven other *Danio* species^8^.

## Main Text References

1 Sucena, E. & Stern, D. L. Divergence of larval morphology between Drosophila sechellia and its sibling species caused by cis-regulatory evolution of ovo/shaven-baby. Proceedings of the National Academy of Sciences of the United States of America 97, 4530–4534, doi: 10.1073/pnas.97.9.4530 (2000).

2 Colosimo, P. F. et al. Widespread parallel evolution in sticklebacks by repeated fixation of Ectodysplasin alleles. Science 307, 1928–1933, doi: 10.1126/science.1107239 (2005).

3 Westerman, E. L. et al. Aristaless Controls Butterfly Wing Color Variation Used in Mimicry and Mate Choice. Curr Biol 28, 3469–3474 e3464, doi: 10.1016/j.cub.2018.08.051 (2018).

4 Barrett, R. D. H. et al. Linking a mutation to survival in wild mice. Science 363, 499–504, doi: 10.1126/science.aav3824 (2019).

5 Hu, Y., Linz, D. M. & Moczek, A. P. Beetle horns evolved from wing serial homologs. Science 366, 1004–1007, doi: 10.1126/science.aaw2980 (2019).

6 Orteu, A. & Jiggins, C. D. The genomics of coloration provides insights into adaptive evolution. Nat Rev Genet, doi: 10.1038/s41576-020-0234-z (2020).

7 Roberts, T. R. The ‘celestial pearl danio’, a new genus and species of colourful minute cyprinid fish from Myanmar (Pisces: Cypriniformes). Raffles Bulletin of Zoology 55, 131–140 (2007).

8 Kullander, S. O. & Fang, F. Danio aesculapii, a new species of danio from south-western Myanmar (Teleostei: Cyprinidae). Zootaxa 2164, 41–48 (2009).

9 Kullander, S. O. & Fang, F. Danio tinwini, a new species of spotted danio from northern Myanmar (Teleostei: Cyprinidae). Ichthyological Exploration of Freshwaters 20, 223–228 (2009).

10 McCluskey, B. M. & Postlethwait, J. H. Phylogeny of zebrafish, a “model species,” within Danio, a “model genus”. Mol Biol Evol 32, 635–652, doi: 10.1093/molbev/msu325 (2015).

11 Frohnhöfer, H. G., Krauss, J., Maischein, H. M. & Nüsslein-Volhard, C. Iridophores and their interactions with other chromatophores are required for stripe formation in zebrafish. Development 140, 2997–3007, doi: 10.1242/dev.096719 (2013).

12 Patterson, L. B. & Parichy, D. M. Interactions with iridophores and the tissue environment required for patterning melanophores and xanthophores during zebrafish adult pigment stripe formation. PLoS genetics 9, e1003561, doi: 10.1371/journal.pgen.1003561 (2013).

13 Singh, A. P., Schach, U. & Nüsslein-Volhard, C. Proliferation, dispersal and patterned aggregation of iridophores in the skin prefigure striped colouration of zebrafish. Nature cell biology 16, 607–614, doi: 10.1038/ncb2955 (2014).

14 Irion, U., Singh, A. P. & Nüsslein-Volhard, C. in Current Topics in Developmental Biology Ch. 8, (Elsevier Inc., 2016).

15 Jinek, M. et al. A programmable dual-RNA-guided DNA endonuclease in adaptive bacterial immunity. Science 337, 816–821, doi: 10.1126/science.1225829 (2012).

16 Hwang, W. Y. et al. Heritable and precise zebrafish genome editing using a CRISPR-Cas system. PloS one 8, e68708, doi: 10.1371/journal.pone.0068708 (2013).

17 Irion, U., Krauss, J. & Nüsslein-Volhard, C. Precise and efficient genome editing in zebrafish using the CRISPR/Cas9 system. Development 141, 4827–4830, doi: 10.1242/dev.115584 (2014).

18 Stern, D. L. Identification of loci that cause phenotypic variation in diverse species with the reciprocal hemizygosity test. Trends Genet 30, 547–554, doi: 10.1016/j.tig.2014.09.006 (2014).

19 Parichy, D. M. & Johnson, S. L. Zebrafish hybrids suggest genetic mechanisms for pigment pattern diversification in Danio. Development genes and evolution 211, 319–328 (2001).

20 Iwashita, M. et al. Pigment pattern in jaguar/obelix zebrafish is caused by a Kir7.1 mutation: implications for the regulation of melanosome movement. PLoS genetics 2, e197, doi: 10.1371/journal.pgen.0020197 (2006).

21 Darwin, C. The Descent of Man, and Selection in Relation to Sex. John Murray, London. (1871).

22 Protas, M. E. & Patel, N. H. Evolution of coloration patterns. Annu Rev Cell Dev Biol 24, 425–446, doi: 10.1146/annurev.cellbio.24.110707.175302 (2008).

23 Cuthill, I. C. et al. The biology of color. Science 357, doi: 10.1126/science.aan0221 (2017).

24 Nüsslein-Volhard, C., Grutzmacher, S. & Howard, J. Animal beauty : on the evolution of biological aesthetics. (The MIT Press, 2019).

25 Parichy, D. M. Evolution of danio pigment pattern development. Heredity 97, 200–210, doi: 10.1038/sj.hdy.6800867 (2006).

26 Kondo, S., Iwashita, M. & Yamaguchi, M. How animals get their skin patterns: fish pigment pattern as a live Turing wave. Int J Dev Biol 53, 851–856, doi: 10.1387/ijdb.072502sk (2009).

27 Singh, A. P. & Nüsslein-Volhard, C. Zebrafish stripes as a model for vertebrate colour pattern formation. Curr Biol 25, R81–92, doi: 10.1016/j.cub.2014.11.013 (2015).

28 Irion, U. & Nüsslein-Volhard, C. The identification of genes involved in the evolution of color patterns in fish. Curr Opin Genet Dev 57, 31–38, doi: 10.1016/j.gde.2019.07.002 (2019).

29 Patterson, L. B. & Parichy, D. M. Zebrafish Pigment Pattern Formation: Insights into the Development and Evolution of Adult Form. Annu Rev Genet 53, 505–530, doi: 10.1146/annurev-genet-112618-043741 (2019).

30 Hirata, M., Nakamura, K., Kanemaru, T., Shibata, Y. & Kondo, S. Pigment cell organization in the hypodermis of zebrafish. Developmental dynamics : an official publication of the American Association of Anatomists 227, 497–503, doi: 10.1002/dvdy.10334 (2003).

31 Hirata, M., Nakamura, K. & Kondo, S. Pigment cell distributions in different tissues of the zebrafish, with special reference to the striped pigment pattern. Developmental dynamics : an official publication of the American Association of Anatomists 234, 293–300, doi: 10.1002/dvdy.20513 (2005).

32 Mahalwar, P., Walderich, B., Singh, A. P. & Nüsslein-Volhard, C. Local reorganization of xanthophores fine-tunes and colors the striped pattern of zebrafish. Science 345, 1362–1364, doi: 10.1126/science.1254837 (2014).

33 Budi, E. H., Patterson, L. B. & Parichy, D. M. Post-embryonic nerve-associated precursors to adult pigment cells: genetic requirements and dynamics of morphogenesis and differentiation. PLoS genetics 7, e1002044, doi: 10.1371/journal.pgen.1002044 (2011).

34 Mongera, A. et al. Genetic lineage labeling in zebrafish uncovers novel neural crest contributions to the head, including gill pillar cells. Development 140, 916–925, doi: 10.1242/dev.091066 (2013).

35 Dooley, C. M., Mongera, A., Walderich, B. & Nusslein-Volhard, C. On the embryonic origin of adult melanophores: the role of ErbB and Kit signalling in establishing melanophore stem cells in zebrafish. Development 140, 1003–1013, doi: 10.1242/dev.087007 (2013).

36 Patterson, L. B., Bain, E. J. & Parichy, D. M. Pigment cell interactions and differential xanthophore recruitment underlying zebrafish stripe reiteration and Danio pattern evolution. Nature communications 5, 5299, doi: 10.1038/ncomms6299 (2014).

37 Singh, A. P. et al. Pigment Cell Progenitors in Zebrafish Remain Multipotent through Metamorphosis. Dev Cell 38, 316–330, doi: 10.1016/j.devcel.2016.06.020 (2016).

38 Parichy, D. M. et al. Mutational analysis of endothelin receptor b1 (rose) during neural crest and pigment pattern development in the zebrafish Danio rerio. Developmental biology 227, 294–306, doi: 10.1006/dbio.2000.9899 (2000).

39 Parichy, D. M. & Turner, J. M. Temporal and cellular requirements for Fms signaling during zebrafish adult pigment pattern development. Development 130, 817–833 (2003).

40 Walderich, B., Singh, A. P., Mahalwar, P. & Nusslein-Volhard, C. Homotypic cell competition regulates proliferation and tiling of zebrafish pigment cells during colour pattern formation. Nat Commun 7, 11462, doi: 10.1038/ncomms11462 (2016).

41 Haffter, P. et al. Mutations affecting pigmentation and shape of the adult zebrafish. Development genes and evolution 206, 260–276, doi: DOI 10.1007/s004270050051 (1996).

42 Eom, D. S. et al. Melanophore migration and survival during zebrafish adult pigment stripe development require the immunoglobulin superfamily adhesion molecule Igsf11. PLoS genetics 8, e1002899, doi: 10.1371/journal.pgen.1002899 (2012).

43 Inaba, M., Yamanaka, H. & Kondo, S. Pigment pattern formation by contact-dependent depolarization. Science 335, 677, doi: 10.1126/science.1212821 (2012).

44 Watanabe, M., Sawada, R., Aramaki, T., Skerrett, I. M. & Kondo, S. The Physiological Characterization of Connexin41.8 and Connexin39.4, Which Are Involved in the Striped Pattern Formation of Zebrafish. J Biol Chem 291, 1053–1063, doi: 10.1074/jbc.M115.673129 (2016).

45 Parichy, D. M. Advancing biology through a deeper understanding of zebrafish ecology and evolution. Elife 4, doi: 10.7554/eLife.05635 (2015).

46 Irion, U. et al. Gap junctions composed of connexins 41.8 and 39.4 are essential for colour pattern formation in zebrafish. Elife 3, e05125, doi: 10.7554/eLife.05125 (2014).

47 Watanabe, M. et al. Spot pattern of leopard Danio is caused by mutation in the zebrafish connexin41.8 gene. EMBO reports 7, 893–897, doi: 10.1038/sj.embor.7400757 (2006).

48 Maderspacher, F. & Nüsslein-Volhard, C. Formation of the adult pigment pattern in zebrafish requires leopard and obelix dependent cell interactions. Development 130, 3447–3457 (2003).

49 Henke, K. et al. Genetic Screen for Postembryonic Development in the Zebrafish (Danio rerio): Dominant Mutations Affecting Adult Form. Genetics 207, 609–623, doi: 10.1534/genetics.117.300187 (2017).

50 Volkening, A. & Sandstede, B. Iridophores as a source of robustness in zebrafish stripes and variability in Danio patterns. Nat Commun 9, 3231, doi: 10.1038/s41467-018-05629-z (2018).

51 Dahal, G. R. et al. An inwardly rectifying K+ channel is required for patterning. Development 139, 3653–3664, doi: 10.1242/dev.078592 (2012).

52 Perathoner, S. et al. Bioelectric signaling regulates size in zebrafish fins. PLoS genetics 10, e1004080, doi: 10.1371/journal.pgen.1004080 (2014).

53 Stewart, S. et al. <em>longfin</em> causes <em>cis</em>-ectopic expression of the <em>kcnh2a ether-a-go-go</em> K^+^ channel to autonomously prolong fin outgrowth. bioRxiv, 790329, doi: 10.1101/790329 (2019).

54 Lee, M. M., Ritter, R., 3rd, Hirose, T., Vu, C. D. & Edwards, A. O. Snowflake vitreoretinal degeneration: follow-up of the original family. Ophthalmology 110, 2418–2426, doi: 10.1016/S0161-6420(03)00828-5 (2003).

55 Hejtmancik, J. F. et al. Mutations in KCNJ13 cause autosomal-dominant snowflake vitreoretinal degeneration. Am J Hum Genet 82, 174–180, doi: 10.1016/j.ajhg.2007.08.002 (2008).

56 Sergouniotis, P. I. et al. Recessive mutations in KCNJ13, encoding an inwardly rectifying potassium channel subunit, cause leber congenital amaurosis. Am J Hum Genet 89, 183–190, doi: 10.1016/j.ajhg.2011.06.002 (2011).

57 Khan, A. O., Bergmann, C., Neuhaus, C. & Bolz, H. J. A distinct vitreo-retinal dystrophy with early-onset cataract from recessive KCNJ13 mutations. Ophthalmic Genet 36, 79–84, doi: 10.3109/13816810.2014.985846 (2015).

58 Pattnaik, B. R. et al. A Novel KCNJ13 Nonsense Mutation and Loss of Kir7.1 Channel Function Causes Leber Congenital Amaurosis (LCA16). Hum Mutat 36, 720–727, doi: 10.1002/humu.22807 (2015).

59 Perez-Roustit, S. et al. Leber Congenital Amaurosis with Large Retinal Pigment Clumps Caused by Compound Heterozygous Mutations in Kcnj13. Retin Cases Brief Rep 11, 221–226, doi: 10.1097/ICB.0000000000000326 (2017).

60 Toms, M. et al. Missense variants in the conserved transmembrane M2 protein domain of KCNJ13 associated with retinovascular changes in humans and zebrafish. Exp Eye Res 189, 107852, doi: 10.1016/j.exer.2019.107852 (2019).

61 Gavrilets, S. & Losos, J. B. Adaptive radiation: contrasting theory with data. Science 323, 732–737, doi: 10.1126/science.1157966 (2009).

62 Spiewak, J. E. et al. Evolution of Endothelin signaling and diversification of adult pigment pattern in Danio fishes. PLoS genetics 14, e1007538, doi: 10.1371/journal.pgen.1007538 (2018).

## Methods References

1 Brand, M., Granato, M. & Nüsslein-Volhard, C. in Zebrafish: A Practical Approach (eds C. Nüsslein-Volhard & R. Dahm) (Oxford University Press, 2002).

2 Lister, J. A., Robertson, C. P., Lepage, T., Johnson, S. L. & Raible, D. W. nacre encodes a zebrafish microphthalmia-related protein that regulates neural-crest-derived pigment cell fate. Development 126, 3757–3767 (1999).

3 Odenthal, J. et al. Mutations affecting xanthophore pigmentation in the zebrafish, Danio rerio. Development 123, 391–398 (1996).

4 Krauss, J., Astrinidis, P., Fröhnhofer, H. G., Walderich, B. & Nüsslein-Volhard, C. transparent, a gene affecting stripe formation in Zebrafish, encodes the mitochondrial protein Mpv17 that is required for iridophore survival. Biol Open 2, 703–710, doi: 10.1242/bio.20135132 (2013).

5 Haffter, P. et al. Mutations affecting pigmentation and shape of the adult zebrafish. Development genes and evolution 206, 260–276, doi: DOI 10.1007/s004270050051 (1996).

6 Watanabe, M. et al. Spot pattern of leopard Danio is caused by mutation in the zebrafish connexin41.8 gene. EMBO reports 7, 893–897, doi: 10.1038/sj.embor.7400757 (2006).

7 Irion, U. et al. Gap junctions composed of connexins 41.8 and 39.4 are essential for colour pattern formation in zebrafish. Elife 3, e05125, doi: 10.7554/eLife.05125 (2014).

8 Parichy, D. M. & Johnson, S. L. Zebrafish hybrids suggest genetic mechanisms for pigment pattern diversification in Danio. Development genes and evolution 211, 319–328 (2001).

9 Parichy, D. M., Elizondo, M. R., Mills, M. G., Gordon, T. N. & Engeszer, R. E. Normal table of postembryonic zebrafish development: staging by externally visible anatomy of the living fish. Developmental dynamics : an official publication of the American Association of Anatomists 238, 2975–3015, doi: 10.1002/dvdy.22113 (2009).

10 Irion, U., Krauss, J. & Nüsslein-Volhard, C. Precise and efficient genome editing in zebrafish using the CRISPR/Cas9 system. Development 141, 4827–4830, doi: 10.1242/dev.115584 (2014).

11 Meeker, N. D., Hutchinson, S. A., Ho, L. & Trede, N. S. Method for isolation of PCR-ready genomic DNA from zebrafish tissues. Biotechniques 43, 610, 612, 614, doi: 10.2144/000112619 (2007).

12 Schindelin, J. et al. Fiji: an open-source platform for biological-image analysis. Nat Methods 9, 676–682, doi: 10.1038/nmeth.2019 (2012).

13 Dobin, A. & Gingeras, T. R. Mapping RNA-seq Reads with STAR. Curr Protoc Bioinformatics 51, 11 14 11–11 14 19, doi: 10.1002/0471250953.bi1114s51 (2015).

14 Love, M. I., Huber, W. & Anders, S. Moderated estimation of fold change and dispersion for RNA-seq data with DESeq2. Genome Biol 15, 550, doi: 10.1186/s13059-014-0550-8 (2014).

15 Bowen, M. E., Henke, K., Siegfried, K. R., Warman, M. L. & Harris, M. P. Efficient mapping and cloning of mutations in zebrafish by low-coverage whole-genome sequencing. Genetics 190, 1017–1024, doi: 10.1534/genetics.111.136069 (2012).

16 Dooley, C. M. et al. Multi-allelic phenotyping--a systematic approach for the simultaneous analysis of multiple induced mutations. Methods 62, 197–206, doi: 10.1016/j.ymeth.2013.04.013 (2013).

17 Poplin, R. et al. Scaling accurate genetic variant discovery to tens of thousands of samples. bioRxiv, 201178, doi: 10.1101/201178 (2018).

18 Li, H. et al. The Sequence Alignment/Map format and SAMtools. Bioinformatics 25, 2078–2079, doi: 10.1093/bioinformatics/btp352 (2009).

19 Notredame, C., Higgins, D. G. & Heringa, J. T-Coffee: A novel method for fast and accurate multiple sequence alignment. J Mol Biol 302, 205–217, doi: 10.1006/jmbi.2000.4042 (2000).

## Figure References

1 McCluskey, B. M. & Postlethwait, J. H. Phylogeny of zebrafish, a “model species,” within Danio, a “model genus”. Mol Biol Evol 32, 635–652, doi: 10.1093/molbev/msu325 (2015).

2 Parichy, D. M., Elizondo, M. R., Mills, M. G., Gordon, T. N. & Engeszer, R. E. Normal table of postembryonic zebrafish development: staging by externally visible anatomy of the living fish. Developmental dynamics : an official publication of the American Association of Anatomists 238, 2975–3015, doi: 10.1002/dvdy.22113 (2009).

3 Irion, U. et al. Gap junctions composed of connexins 41.8 and 39.4 are essential for colour pattern formation in zebrafish. Elife 3, e05125, doi: 10.7554/eLife.05125 (2014).

4 Haffter, P. et al. Mutations affecting pigmentation and shape of the adult zebrafish. Development genes and evolution 206, 260–276, doi: DOI 10.1007/s004270050051 (1996).

5 Watanabe, M. et al. Spot pattern of leopard Danio is caused by mutation in the zebrafish connexin41.8 gene. EMBO reports 7, 893–897, doi: 10.1038/sj.embor.7400757 (2006).

6 Eom, D. S. et al. Melanophore migration and survival during zebrafish adult pigment stripe development require the immunoglobulin superfamily adhesion molecule Igsf11. PLoS genetics 8, e1002899, doi: 10.1371/journal.pgen.1002899 (2012).

7 Iwashita, M. et al. Pigment pattern in jaguar/obelix zebrafish is caused by a Kir7.1 mutation: implications for the regulation of melanosome movement. PLoS genetics 2, e197, doi: 10.1371/journal.pgen.0020197 (2006).

8 Henke, K. et al. Genetic Screen for Postembryonic Development in the Zebrafish (Danio rerio): Dominant Mutations Affecting Adult Form. Genetics 207, 609–623, doi: 10.1534/genetics.117.300187 (2017).

9 Hejtmancik, J. F. et al. Mutations in KCNJ13 cause autosomal-dominant snowflake vitreoretinal degeneration. Am J Hum Genet 82, 174–180, doi: 10.1016/j.ajhg.2007.08.002 (2008).

10 Sergouniotis, P. I. et al. Recessive mutations in KCNJ13, encoding an inwardly rectifying potassium channel subunit, cause leber congenital amaurosis. Am J Hum Genet 89, 183–190, doi: 10.1016/j.ajhg.2011.06.002 (2011).

11 Khan, A. O., Bergmann, C., Neuhaus, C. & Bolz, H. J. A distinct vitreo-retinal dystrophy with early-onset cataract from recessive KCNJ13 mutations. Ophthalmic Genet 36, 79–84, doi: 10.3109/13816810.2014.985846 (2015).

12 Perez-Roustit, S. et al. Leber Congenital Amaurosis with Large Retinal Pigment Clumps Caused by Compound Heterozygous Mutations in Kcnj13. Retin Cases Brief Rep 11, 221–226, doi: 10.1097/ICB.0000000000000326 (2017).

13 Toms, M. et al. Missense variants in the conserved transmembrane M2 protein domain of KCNJ13 associated with retinovascular changes in humans and zebrafish. Exp Eye Res 189, 107852, doi: 10.1016/j.exer.2019.107852 (2019).

## Supplementary Information References

1 Ogino, S. et al. Standard mutation nomenclature in molecular diagnostics: practical and educational challenges. J Mol Diagn 9, 1–6, doi: 10.2353/jmoldx.2007.060081 (2007).

2 Hibino, H. et al. Inwardly rectifying potassium channels: their structure, function, and physiological roles. Physiol Rev 90, 291–366, doi: 10.1152/physrev.00021.2009 (2010).

3 Haffter, P. et al. Mutations affecting pigmentation and shape of the adult zebrafish. Development genes and evolution 206, 260–276, doi:DOI 10.1007/s004270050051 (1996).

4 Irion, U. et al. Gap junctions composed of connexins 41.8 and 39.4 are essential for colour pattern formation in zebrafish. Elife 3, e05125, doi: 10.7554/eLife.05125 (2014).

5 Henke, K. et al. Genetic Screen for Postembryonic Development in the Zebrafish (Danio rerio): Dominant Mutations Affecting Adult Form. Genetics 207, 609–623, doi: 10.1534/genetics.117.300187 (2017).

6 Iwashita, M. et al. Pigment pattern in jaguar/obelix zebrafish is caused by a Kir7.1 mutation: implications for the regulation of melanosome movement. PLoS genetics 2, e197, doi: 10.1371/journal.pgen.0020197 (2006).

7 Maderspacher, F. & Nüsslein-Volhard, C. Formation of the adult pigment pattern in zebrafish requires leopard and obelix dependent cell interactions. Development 130, 3447–3457 (2003).

